# Target of Rapamycin: function in abiotic stress tolerance in Arabidopsis and its involvement in a possible cross-talk with ribosomal proteins

**DOI:** 10.1101/2020.01.15.907899

**Authors:** Achala Bakshi, Mazahar Moin, M. S. Madhav, Meher B. Gayatri, Aramati B. M. Reddy, Raju Datla, P. B. Kirti

**Author notes:** **Correspondence to:** Achala Bakshi, P. B. Kirti. **Contact details of authors:** Mazahar Moin; M. S. Madhav; Meher B. Gayatri; Aramati B. M. Reddy; Raju Datla. **Abbreviations:** *AtTOR*, *Arabidopsis thaliana Target of Rapamycin*; *TOR*-OE, *TOR* overexpressing; RPL, Ribosomal protein large subunit; RPS, Ribosomal protein small subunit; RPs, Ribosomal Proteins; TFs, Transcription factors; WT, Wild Type; DAG, Days after Germination; PEG, Polyethylene glycol; RSK, Ribosomal S6 kinase.

## Abstract

The Target of Rapamycin (TOR) protein kinase reprograms cellular metabolism under various environmental stresses. The overexpression of *TOR* in Arabidopsis resulted in increased plant growth including yield and biomass when compared with the wild type under both controlled and limited water conditions. In the present investigation, we report that Arabidopsis plants overexpressing TOR exhibited enhanced tolerance to the osmotic and salt stress treatments. Further to determine the role of TOR in abiotic stresses other than water limiting conditions, which were observed earlier in rice, we have treated high and medium *TOR* expressing Arabidopsis plants, ATR-1.4.27 and ATR-3.7.32 respectively, with stress-inducing chemical agents such as Mannitol (100 mM), NaCl (150 mM), Sorbitol (200 mM) and PEG (7%). Both the lines, ATR-1.4.27 and ATR-3.7.32 exhibited enhanced tolerance to these stresses. These lines also had increased proline and total chlorophyll contents under stress conditions compared with their corresponding WT counterparts. The upregulation of several osmotic stress inducible genes in Arabidopsis transgenic lines indicated the role of TOR in modulating multi-stress tolerance. In the present investigation, we have also analyzed the transcriptional upregulation of ribosomal protein large and small subunit (RPL and RPS) genes in *AtTOR* overexpressing rice transgenic lines, TR-2.24 and TR-15.1 generated earlier (Bakshi et al., 2017a), which indicated that TOR also positively regulates the transcription of ribosomal proteins (RP) along with the synthesis of rRNAs. Also, the observations from phosphoproteomic analysis in SALK lines of various Arabidopsis T-DNA insertion mutants of ribosomal proteins showed differential regulation in phosphorylation of p70kDa ribosomal protein S6K1 and comparative analysis of phosphorylation sites for RSK (Ribosomal S6 Kinases) in RPL6, RPL18, RPL23, RPL24 and RPS28C proteins of Arabidopsis, Interestingly, rice showed similarity in their peptide sequences and Ser/Thr positions. These results suggest that the phosphorylation of S6K1 is controlled by loss/ inhibition of ribosomal protein function to switch ‘on’/ ‘off’ the translational regulation for balanced growth and the pathways of both RPs and TOR are interlinked in a cyclic manner via phosphorylation of S6K1 as a modulatory step.

## 1. Introduction

Target of Rapamycin (TOR) is a conserved eukaryotic Ser/Thr protein kinase, which controls signaling networks involved in cell growth and development and is a key regulator of cellular metabolism in plants and mammals (Loewith et al. 2002). TOR modulates growth and development in association with the nutrient availability and energy status of the cell. TOR also plays a central role in the regulation of low energy signaling and metabolic reprogramming under stress conditions (Caldana et al. 2013; Tomé et al. 2014). The TOR protein exhibits five conserved domains; these include a HEAT repeat domain, a FAT domain, a FRB domain, a Ser/Thr kinase domain and a FATC domain from N to C-terminus regions, which together play important roles in regulating various functions of the cell. Plants and mammals have single copy of the TOR gene, whereas the yeast TOR is encoded by two TOR genes, TOR1 and TOR2 (Cafferkey et al. 1993). Each one of these domains has specific function; the HEAT domain is involved in rRNA synthesis (Ren et al. 2011), the FRB domain provides the binding site for a macrocyclic, immunosuppressant drug, Rapamycin. The FAT and FATC domains are involved in scaffolding and protein-protein interactions. TOR protein and their interacting partners function as two complexes, TORC1 and TORC2 (Loewith et al. 2002). TORC1 is a complex of TOR protein, LST8 and RAPTOR that regulates basic functions for cell survival such as cell growth, ribosome biogenesis, protein translation in response to nutrients and energy (Kim et al. 2002). The TORC2 complex regulates cytoskeletal structure, actin polarization, cell polarity and the complex consists of TOR protein, LST8, SIN1 and RICTOR (Kunz et al. 2000; Wullschleger et al. 2006).

The ribosome large and small subunits comprise ribosomal proteins and various rRNAs. TOR regulates the rRNA synthesis and hence, ribosome biogenesis in yeast, mammals and plants (Hay and Sonenberg, 2004; Urban et al. 2007; Ren et al. 2011), whereas the regulation of genes encoding large and small subunit RPs is not well understood. In yeast, the transcription of RP genes is regulated by TOR-mediated regulation of Forkhead transcription factor (FHL1) along with its co-activator, IFH1 and the repressor, CRF1 (Martin et al. 2004; Wade et al. 2004). Under favorable conditions, FHL1 binds to RP gene promoter regions and activates their transcription, whereas under unfavorable conditions, the FHL binds to its repressor CRF1, which leads to the suppression of RP gene transcription (Martin et al. 2004).

The Arabidopsis and rice plants constitutively overexpressing *AtTOR* exhibited enhanced growth, increased seed yield, leaf size and improved water-use efficiency (Deprost et al. 2007; Ren et al. 2011; Bakshi et al. 2017). The AtTOR protein has also been linked to regulate root growth in nitrogen starved condition positively (Deprost et al. 2007). The expression levels of TOR have also been correlated with plant growth and productivity in Arabidopsis (Deprost et al. 2007; Ren et al. 2011, Xiong and Sheen, 2012) and rice (Bakshi et al. 2017). The detailed multifunctional roles of the TOR protein have been reviewed recently by Bakshi et al. (2019). TOR negatively regulates autophagy, which is induced under nutrient limiting conditions (Liu and Bassham, 2010). The environmental stresses activate SnRKs, which negatively regulate TOR activity and act antagonistically with respect to the TOR protein (Tomé et al. 2014). The rapamycin or constitutive *TOR* knockdown Arabidopsis lines had increased raffinose and galactinol metabolism, which are generally accumulated under abiotic stress conditions (Dobrenel et al. 2011; Caldana et al. 2013). The TORC1-RAPTOR1-S6K1 signaling regulates responses to abiotic stresses in plants (Mahfouz et al. 2006). These findings and research advances suggest that TOR signaling is an important target for improving stress tolerance and yields in crop plants. Despite these advances, the molecular mechanisms of TOR underpinning the abiotic stress responses are still unclear. In the present study, we explored the effects of overexpression of *TOR* in Arabidopsis in response to various abiotic stress conditions. In the *TOR*-OE Arabidopsis transgenic plants display improved shoot and root and activation of various stress inducible genes when challenged with different abiotic stress conditions. We have campared the performance of the *TOR* overexpression phenotypes under abiotic stress conditions in two plant systems; rice and Arabidopsis. The *AtTOR* expressing rice transgenic lines also exhibited similar tolerance to abiotic stress treatments (Bakshi et al., 2017). Furthermore, the key findings from this study also showed the important involvement of TOR pathway in ribosomal assembly by controlling RP transcription and translation via S6K(1) phosphorylation in plants.

## 2. Material and Methods

### 2.1. *AtTOR* DNA vector

The full-length *TOR* cDNA (7.4 kb) was amplified from *Arabidopsis thaliana* (Col 0) and cloned into pEarleyGate-203 (Ren et al. 2011). The derivative binary vector also carried the *bar* gene under mannopine synthase promoter as a plant selection marker for the herbicide, phosphinothricin (PPT).

### 2.2. Generation of *TOR*-OE lines in *Arabidopsis thaliana* and rice

The wild type (WT) *Arabidopsis thaliana* (Col 0) plants were grown on solid ½ concentration of Murashige and Skoog (MS) medium at 22 ± 2°C following 8h light and 16 h dark photoperiod for vegetative growth. The light intensity used for plant growth was 100-150 μmol m^−2^ s^−1^. Following the vegetative phase of growth, plants were shifted to 16 h light and 8 h dark photoperiod to induce flowering. The pEarlyGate 203 vector carrying 35S:*TOR* cassette was mobilized into *Agrobacterium* EHA105 strain by the standard freeze-thaw method of transformation.

The *Arabidopsis thaliana* plants were transformed using *Agrobacterium-*mediated floral dip method as described by Clough and Bent (1998). The *Agrobacterium* strain carrying the vector was grown in 10 ml Luria Bertini broth (LB) at 28°C for 24 h in an orbital shaker at 200 rpm. Then, 1% of this culture was re-inoculated in 100 ml LB overnight. The culture was then centrifuged and resuspended in suspension medium (half-strength MS salts, 0.005% BAP, 5% Sucrose and 0.2% surfactant, Silwet L-77 and 200 μM Acetosyringone). The immature floral buds were then dipped in *Agrobacterium* suspension for 2 min followed by vacuum infiltration. The *Agrobacterium*-infected plants were covered with transparent plastic films to maintain humid conditions and kept in dark for 24 h. These plants were then transferred to 16 h light /8 h dark conditions and seeds were harvested. The putative independent transformed lines of *TOR*-OE are identified first by selecting the seedlings on herbicide PPT (include the details of PPT source), followed by growth of these in individual pots.

We have earlier reported on the developmenty of rice transgenic lines overexpressing the *AtTOR* gene using an *in planta* transformation protocol in a widely cultivated (BPT-5204) variety of ssp. *indica*, using the same binary vector and the lines were characterized for an agronomically important trait, water use efficiency (Bakshi et al., 2017).

### 2.3. Screening of positive *TOR*-OE transgenic lines

The T_1_ and T_2_ generation seeds obtained from the *Agrobacterium* treated T_0_ Arabidopsis plants were surface sterilized with 4% sodium hypochlorite followed by five stringent washes with sterile double-distilled water and allowed to germinate on solid ½ MS medium containing 10 mg ml^−1^ phosphinothricin (PPT). The transformation efficiency (%) was calculated as percent of seeds germinated on PPT in relation to the total number of seeds inoculated. (PPT resistant seedlings)/ (Total number of seedlings tested) × 100 (Supplementary Fig. 1a, 1b, 2a, 2b, & 2c). Nearly 100% germination of seedlings of *TOR-*OE transgenic lines was observed on selection medium in T_2_ generation indicating the homozygous nature of plants (Supplementary Fig. 1a, 1b, 2a & 2b). The T_1_ and T_2_ transgenic plants were further confirmed by PCR amplification of different elements present within the T-DNA using primers specific to the *bar* gene, CaMV35S promoter and kinase domain of *Arabidopsis thaliana TOR* gene (Supplementary Fig. 1c, Supplementary Table.1).. The high *AtTOR* expressing rice transgenic lines, TR-15.1 and TR-2.24 used in the analysis of RP genes expression were previously generated and screened using a similar method (Bakshi et al. 2017).

Genomic DNA was extracted from T_1_ and T_2_ transgenic plants of Arabidopsis, rice and their corresponding WT plants. About 50-100 ng of genomic DNA was used in 25 µl of PCR reactions containing 2.5 µl 10X buffer, 10 µM of each primer and 1 unit *Taq* polymerase. The 35S: *AtTOR* plasmid was used as a positive control (PC) and WT was used as negative control (NC). PCR was performed with an initial denaturation at 94°C for 4 min followed by 35 repeated cycles of 94°C for 1 min, 56°C for 50 sec, and 72°C for 1 min. The PCR cycle was terminated with a final extension at 72°C for 10 min. Seeds obtained from PCR confirmed T_1_-transgenic plants were collected and processed for germination on solid ½ MS containing PPT (10 mg ml^−1^) to advance to T_2_ generation (Supplementary Fig. 1 & 2, Supplementary Table. 1). The PPT resistant T_3_ generation of Arabidopsis transgenic (homozygous) lines were used for abiotic stress tolerance analysis. The ∑ Chi-Square (χ2) analysis was performed to determine the deviation from the Mendelian segregation in the T_3_-generation of the *TOR*-OE Arabidopsis lines germinated on Basta selection medium (Supplementary Table. 2).

### 2.4. Semi-quantitative (semi-Q) and Quantitative real-time PCR (qRT-PCR) analyses

Total RNA was extracted from the leaves of *TOR*-OE Arabidopsis and fullength *AtTOR* expressing rice transgenic (Bakshi et al., 2017) plants using Tri-Reagent (Takara Bio, UK) following the manufacturer’s protocol. Total RNA was isolated from the leaves of one-month-old Arabidopsis transgenic and wild-type (WT) plants. The RNA pellet was dissolved in nuclease-free water and all the RNA isolation steps were carried out at 4°C and the vials used in the isolation process were treated with DiEthyl PyroCarbonate (DEPC) prior to use. The quality and quantity of extracted RNA was checked on 1.2% agarose gel prepared in TBE (Tris-borate-EDTA) buffer and quantified using a nano-spectrophotometer. Total RNA (2 μg) was used to synthesize the first strand cDNA using SMART^TM^ MMLV Reverse Transcriptase (Takara Bio, USA). The synthesized cDNA was diluted seven times (1:7) and 2 µl of this dilution was used for analyzing the transcript level of *TOR* gene using primers that specifically bind and amplify Arabidopsis *TOR* gene. The *Actin* and *Tubulin* genes were used as endogenous reference genes for normalization in qRT-PCR analyses and Actin was used for semi-Q analysis and for separation of both Arabidopsis and rice lines. The T_3_-generation of *TOR*-OE Arabidopsis transgenic plants were separated into low, medium and high expression lines of *TOR* (Fig. 1a, 1b, 1c & 1d). The Arabidopsis WT cDNA was used as positive control and PCR mix without template was used as negative control for semiQ PCR of *TOR*-OE lines. The conditions used in semi-Q PCR include an initial denaturation at 94°C for 3 min followed by 26 repeated cycles of 94°C for 30 sec, 61.9°C for 25 sec and 72°C for 30 sec with a final extension for 5 min at 72°C. The band intensity of products obtained from semi-Q PCR was observed on the agarose gel and was further characterized with qRT-PCR using SYBR Green ® Premix (Takara Bio, USA). Specific primers were designed for studying expression of stress related genes in Arabidopsis and RP genes in rice using primer3 (v.0.4.0) online tool. The transcript level of stress inducible genes in high expression *TOR*-OE Arabidopsis lines were analyzed by qRT-PCR. List of primers used in PCR, SemiQ PCR and qRT-PCR are detailed in Supplementary Tables, 1, 3, 4 & 5. The qRT-PCR data was analyzed with three biological and three technical replicates according to the ΔΔC_T_ method (Livak and Schmittgen, 2001). The *Actin1* and β-*tubulin* genes were used for rice qRT-PCR analysis of RP genes and *Actin2* and α-*tubulin* genes were used for normalization of Arabidopsis specific stress inducible genes. Similar semi-Q and qRT-PCR analyses were performed to separate *AtTOR* expressing rice transgenic plants and the high expression lines were used for the RP genes expression study (Bakshi et al. 2017). Also, similar qRT-PCR method was used to analyse transcript levels of RP genes in high *AtTOR* expressing rice lines

**Figure 1.**
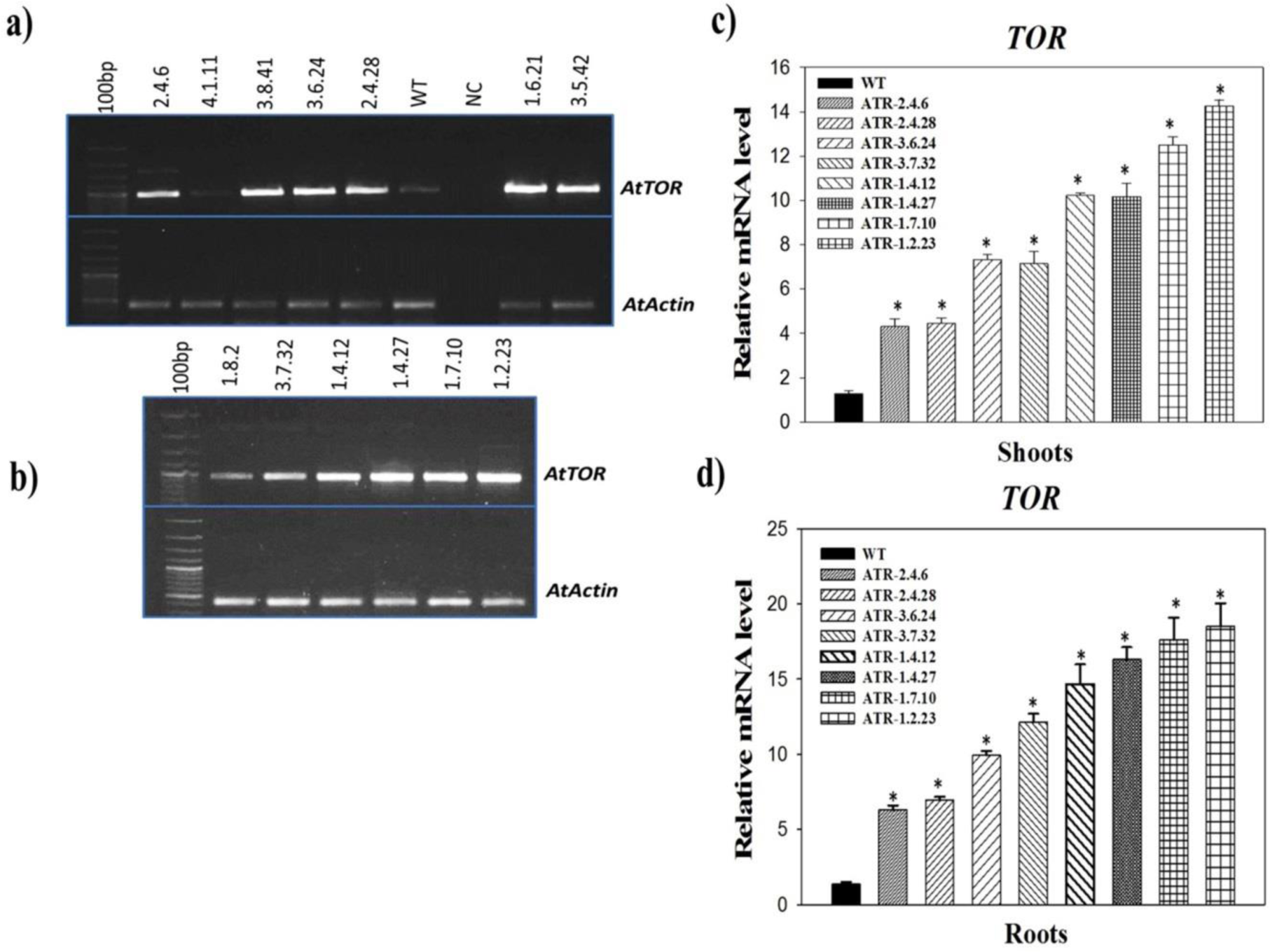
Semi-Q and qRT-PCR analysis of *TOR*-OE Arabidopsis transgenic plants. (a and b) The Arabidopsis actin (*Act2*) was used as an internal reference gene. The AtTOR specific kinase was used to assess the transcript levels in TOR-OE lines. Transgenic lines, ATR-1.4.12, ATR-1.4.27, ATR-1.7.10, ATR-1.2.23, ATR-1.6.21, ATR-3.5.42 and ATR-3.8.41 had high intensity bands. The lines, ATR-3.7.32, ATR-3.6.24 ATR-2.4.6 and ATR-2.4.28 had medium intense bands, whereas lines, ATR-1.8.2 and ATR-4.1.11 had low band intensities on agarose gel.The transcript level of *AtTOR* was analyzed using qRT-PCR. (a) The expression analysis of TOR gene in shoots of T_3_ generation transgenic plants. (b) Graphical representation of expression analysis of TOR in roots of transgenic lines. Arabidopsis actin (*Act2*) was used as an internal reference gene for normalization.

### 2.5 Abiotic Stress treatments

The *TOR*-OE transgenic plants in T_3_ generation of Arabidopsis were analyzed for tolerance to various abiotic stresses. The seeds of transgenic and WT plants were surface sterilized with 4% sodium hypochlorite followed by five stringent washes with sterile double distilled water and were germinated on solid half strength Murashige and Skoog (MS) medium for 15d (Murashige and Skoog, 1962). The plants after 15 days after germination (DAG) were transferred to stress media containing half-MS salts and stress inducing supplements such as sodium chloride (NaCl, 150 mM), polyethylene glycol (PEG-8000, 7%), sorbitol (200 mM) and Mannitol (100 mM). Differences in growth parameters of transgenic and WT plants were observed after one week of each one of the treatments (Fig. 2a, 2b, 2c & 2d). The transgenic and WT plants were also grown on half-MS without any stress supplement that served as untreated control. The root length, fresh weight of 10 plants, chlorophyll and proline contents of transgenic and WT plants were observed. In our earlier report, the two high *AtTOR* expressing rice transgenic lines TR-2.24 and TR-15.1 were treated with osmotic (15% w/v, PEG-8000) and salt (NaCl, 300 mM) stress and the transcript levels of rice specific stress inducible genes were analyzed (Bakshi et al., 2017).

**Figure 2.**
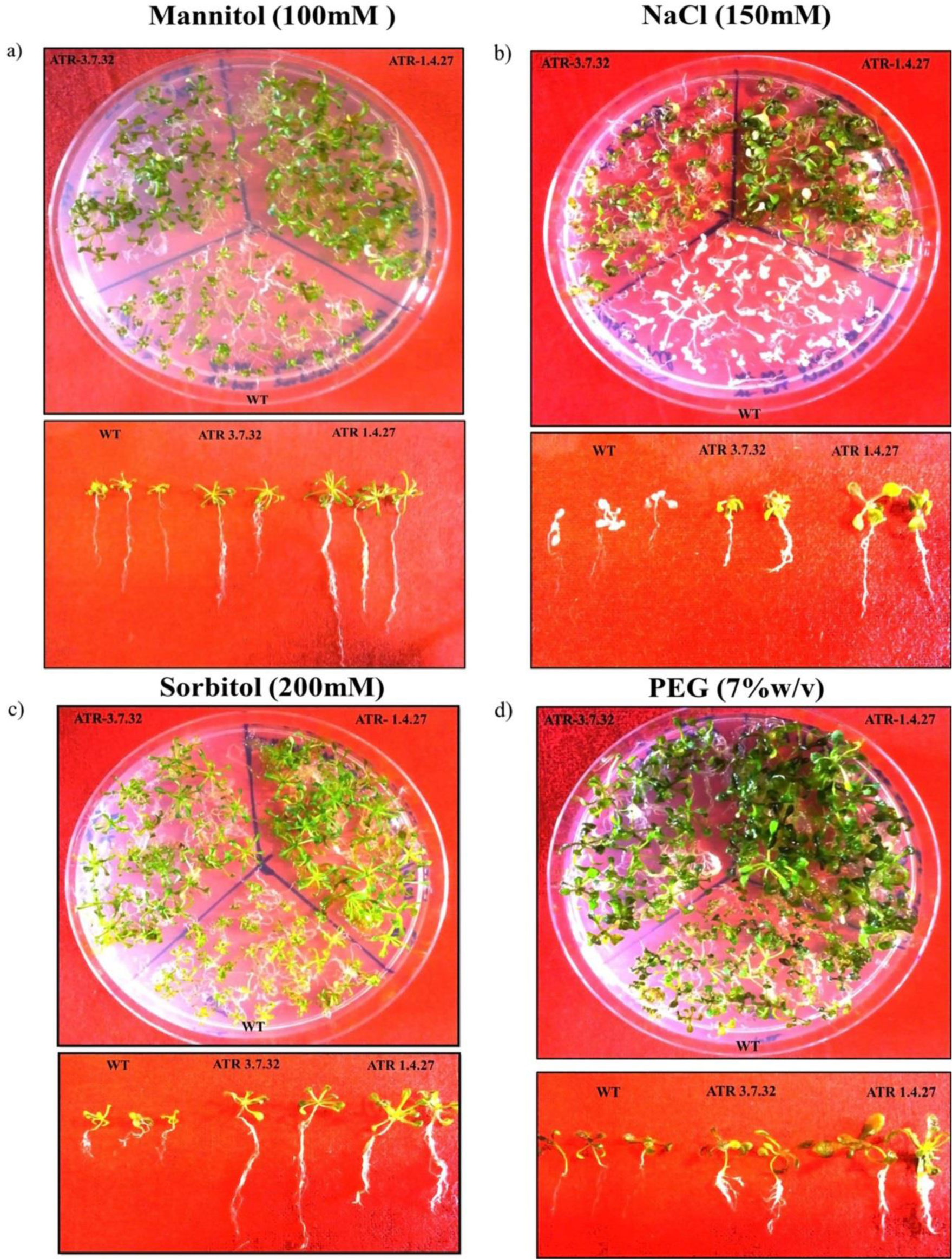
Seedling assay of *TOR*-OE Arabidopsis lines. The 15 DAG high and medium *TOR* transgenic lines, ATR-1.4.27 and ATR-3.7.32, respectively along with the WT were treated with (a) Mannitol (100 mM) (b) NaCl (150mM), (c) Sorbitol (200mM), (d) PEG (7%w/v). The shoot and root growth were observed after 7DAG on treatment of transgenic lines and WT.

The qRT-PCR was performed to analyze the transcript levels of different stress inducible genes using SYBR Green® Premix (Takara Bio, USA) as described earlier. The expression analysis in various treatments was performed using qRT-PCR as described earlier (Supplementary Table. 5). The expression analysis was performed with three biological and three technical replicates and data with the statistically significant differences (P < 0.05) was represented with asterisks (*) in the figures. The expression data of stress induced genes was also statistically tested with one-way ANOVA and represented with a Holm-Bonferroni correction of p-values at P < 0.05 (Supplementary Table. 6; Holm, 1979). The statistical significance was represented with asterisks in bar diagrams at P<0.05 using SigmaPlot v.11.

### 2.7. Estimation of chlorophyll contents

The chlorophyll a, b and total chlorophyll contents in response to treated and untreated conditions in WT and *TOR*-OE Arabidopsis transgenic lines were estimated as described by Hiscox and Israelstam (1979). About 100 mg of leaf tissue was ground to fine powder in DMSO and absorption of the extracts was measured at OD_663_ nm and OD_645_ nm using a UV-Spectrophotometer. The chlorophyll-a, chlorophyll-b and total chlorophyll contents were then calculated as described by Arnon, (1949).

### 2.8. Proline estimation in treated *TOR*-OE Arabidopsis lines

The treated and untreated plants were used for proline estimation following a method described by Bates et al. (1973). About 100 mg plants of each transgenic and WT was homogenized in 5 ml of 3% sulfosalicylic acid and the homogenate was then centrifuged at a speed of 12000 rpm for 15 min. The aqueous phase (0.4 ml) was extracted into a fresh 2 ml tube. Then, equal volumes of acid ninhydrin (prepared in 6N Ortho phosphoric acid) and glacial acetic acid were added to the supernatant and incubated at 100°C. After 1 h of incubation, the tubes were transferred to ice to terminate the reaction. The reaction mixture was extracted with 0.8 ml Toluene and the absorbance was measured at 520 nm wavelength using Toluene as a blank. Proline dilutions were used to prepare a standard curve and proline content in samples was calculated from the standard curve as described by Bates et al. (1973).

### 2.9. Nucleotide Sequence Retrieval of rice RPS and RPL genes

The sequences of ribosomal protein large and small subunit genes of rice (RPL and RPS) were retrieved from RGAP-DB. The sequences were validated in RAP-DB, NCBI, and various other databases to ensure that the sequences were gene specific as described by Moin et al. (2016, 2017) and Saha et al. (2017) (Supplementary Table. 3 & 4).

### 2.10. Retrieval of RPL and RPS protein sequences and identification of Ser/Thr phosphorylation sites in Arabidopsis and Rice

The p70kDa, Ribosomal Protein Small Subunit 6-kinase 1 (RPS6K1) is a Ser/ Thr protein kinase of type B of the AGC kinase family, which phosphorylates RPS6 at several Serine and Threonine residues for the translational initiation. To identify the similar phosphorylation sites at Ser/Thr residues, the Arabidopsis and rice RPS6, RPL6, RPL18, RPL23, RPL24 and RPS28 protein sequences were retrieved from NCBI, UNIPROT and Ensemble Plants and the retrieved sequences were validated in TAIR and RAPDB databases. The obtained sequences were analyzed for the presence of Ser/ Thr phosphorylation sites by PKA, PKB, PKC, PKG and RSK protein kinases of AGC kinase family using NetPhos 3.1 Server (Table. 1; http://www.cbs.dtu.dk/services/NetPhos/). The multiple sequence alignment was performed using Clustal Omega (Fig. 11; https://www.ebi.ac.uk/Tools/msa/clustalo/) with the selected RPL and RPS proteins to check similarity between various Serine/Threonine phosphorylation sites (Table 1; Fig. 11a, b, c, d, e, f)

**Table 1.**
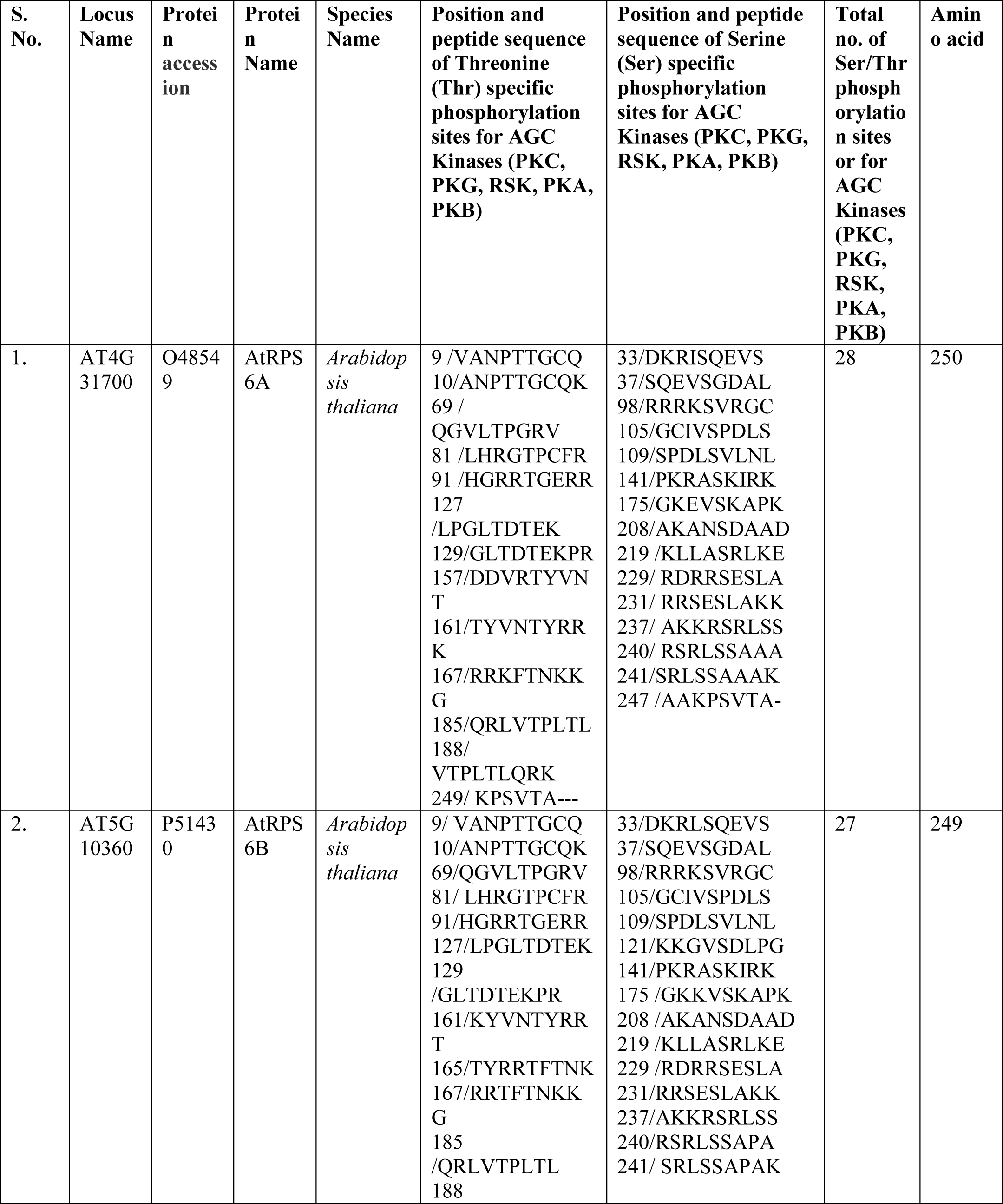
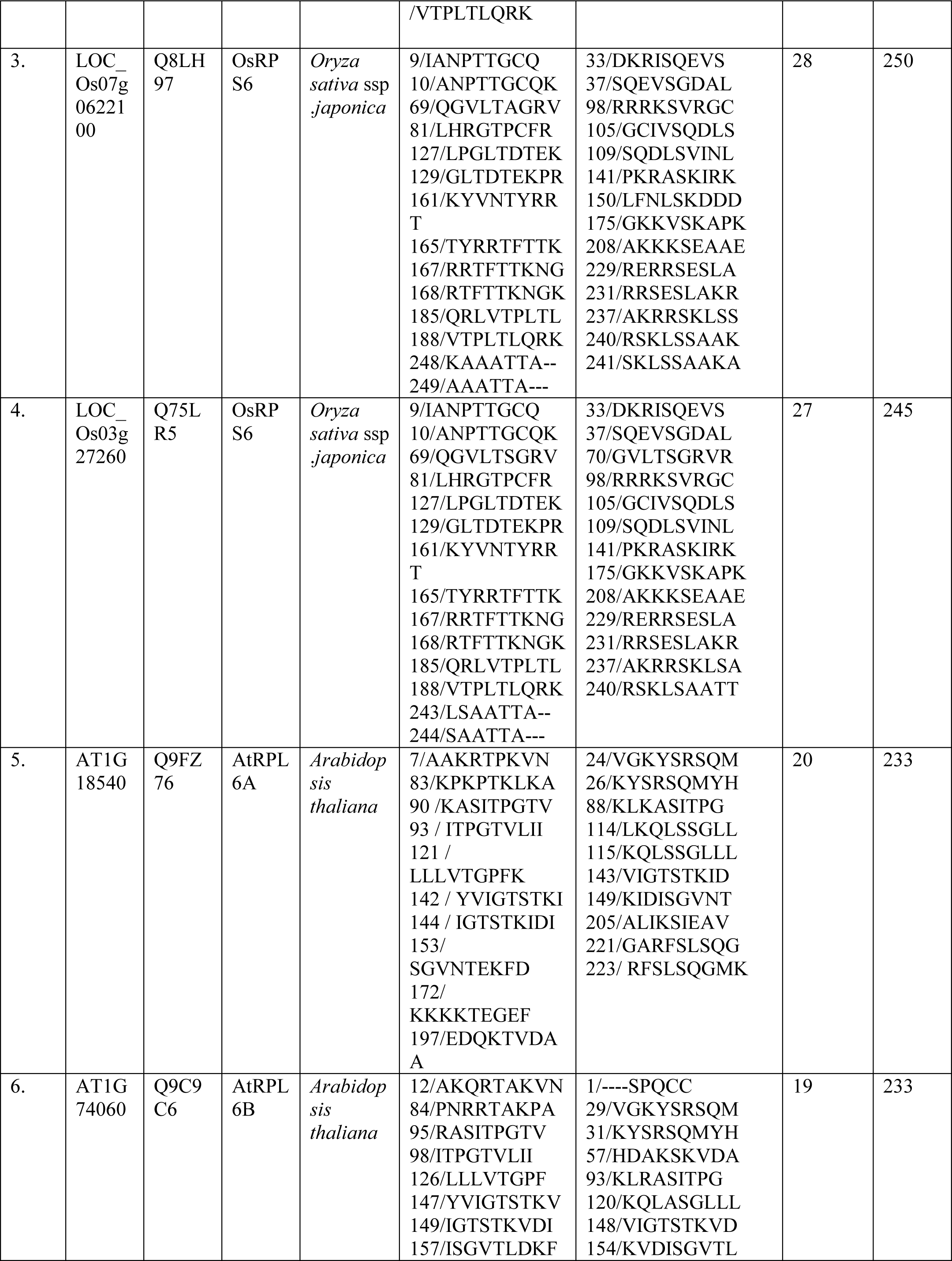
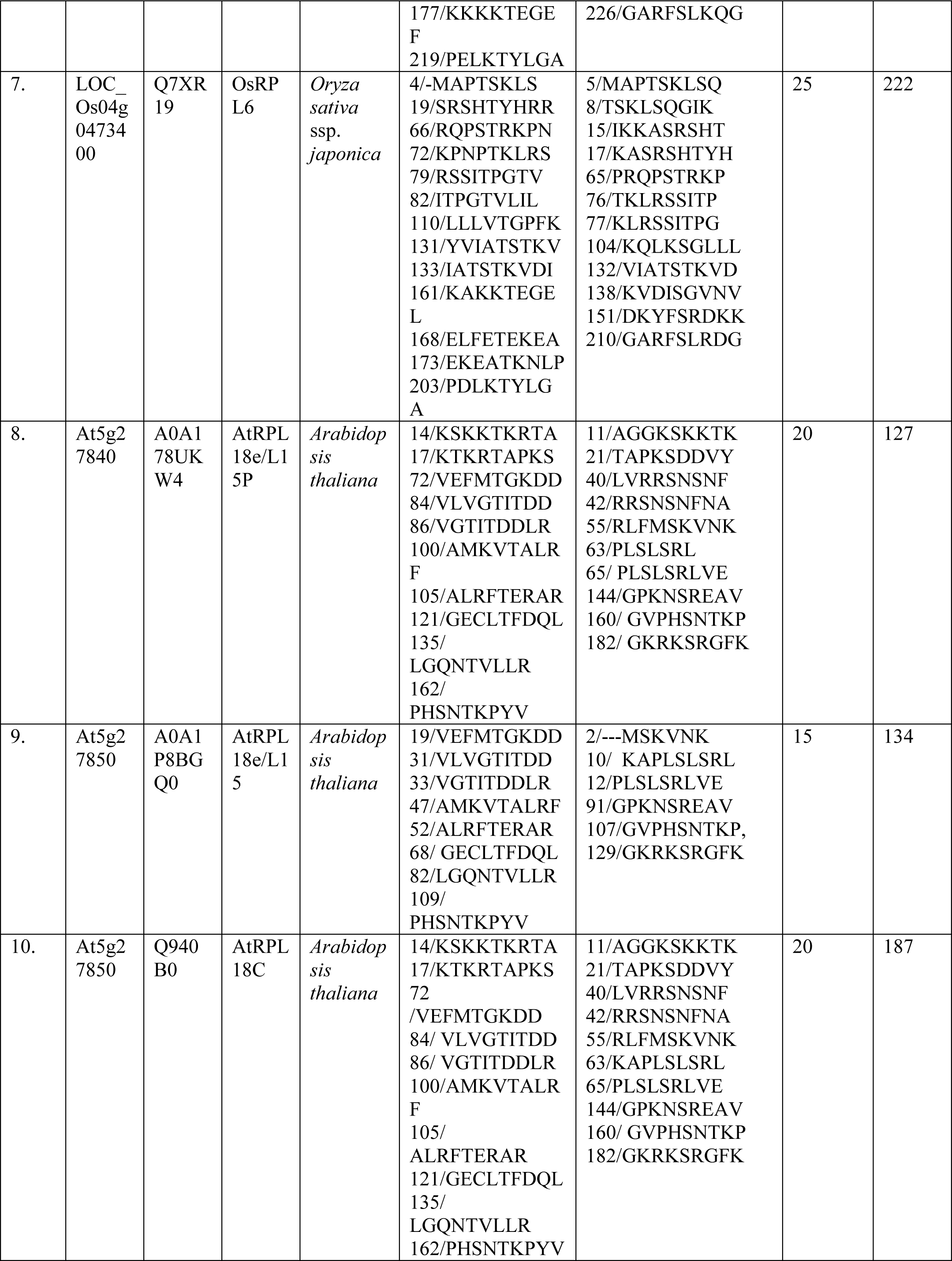
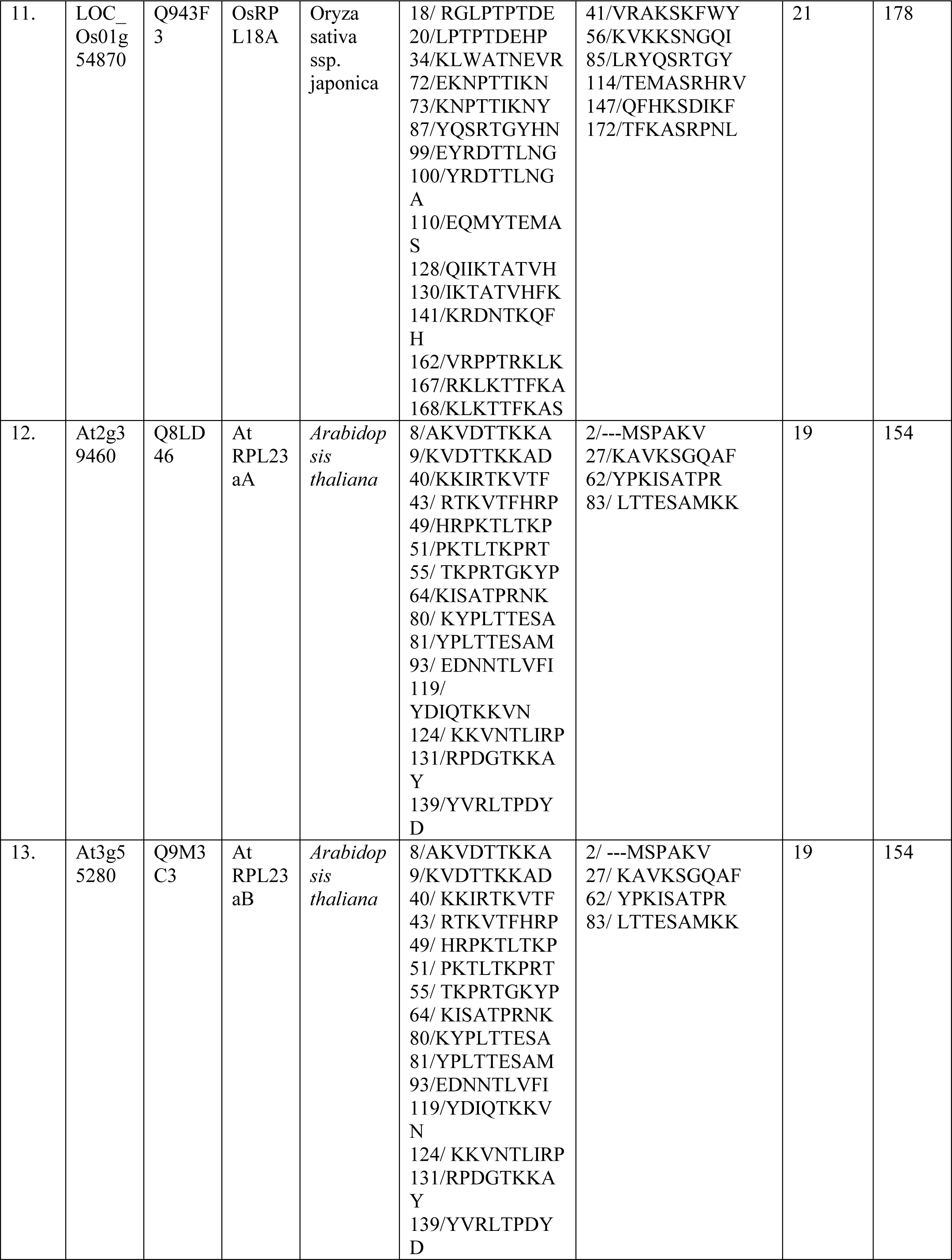
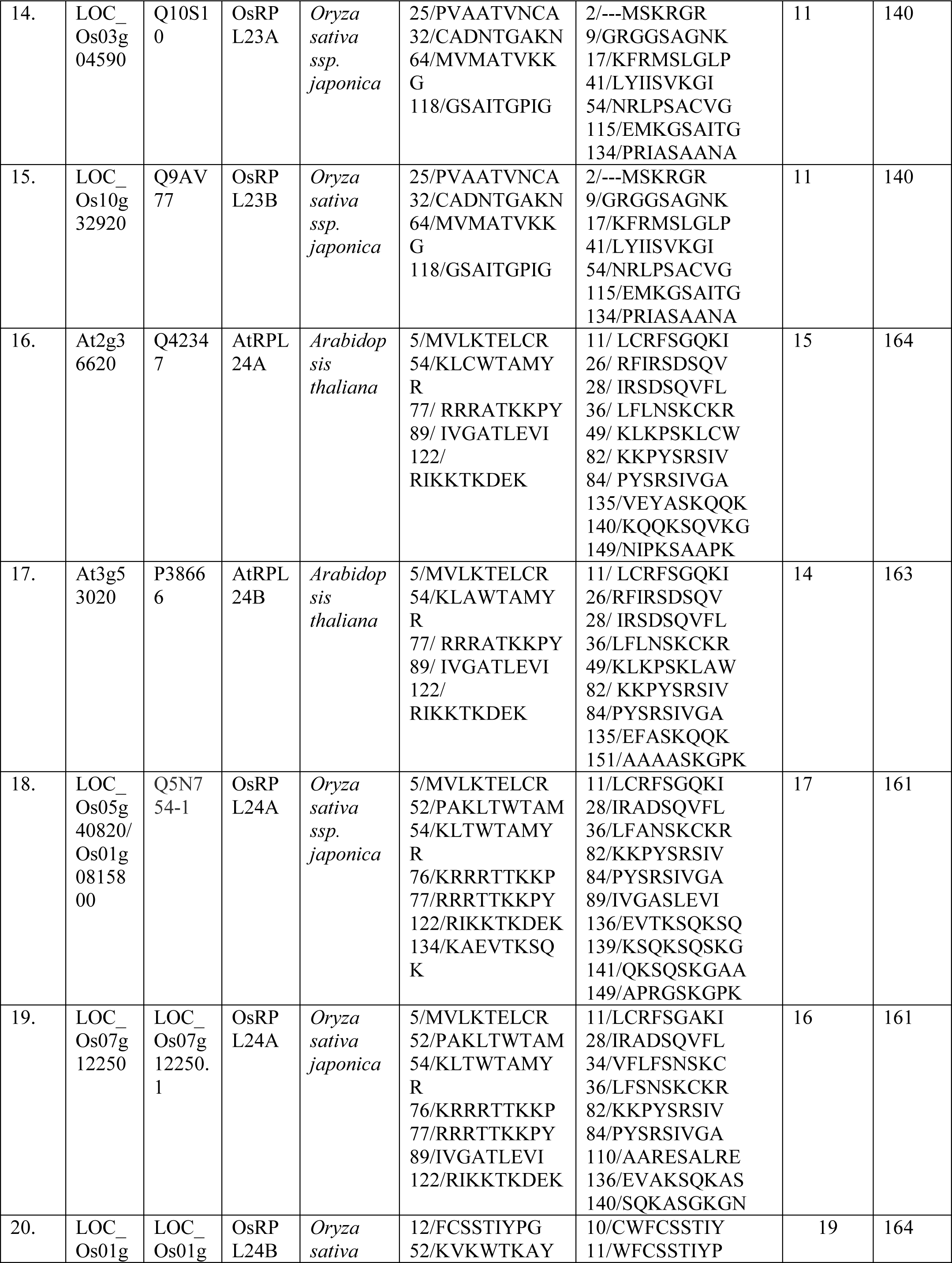
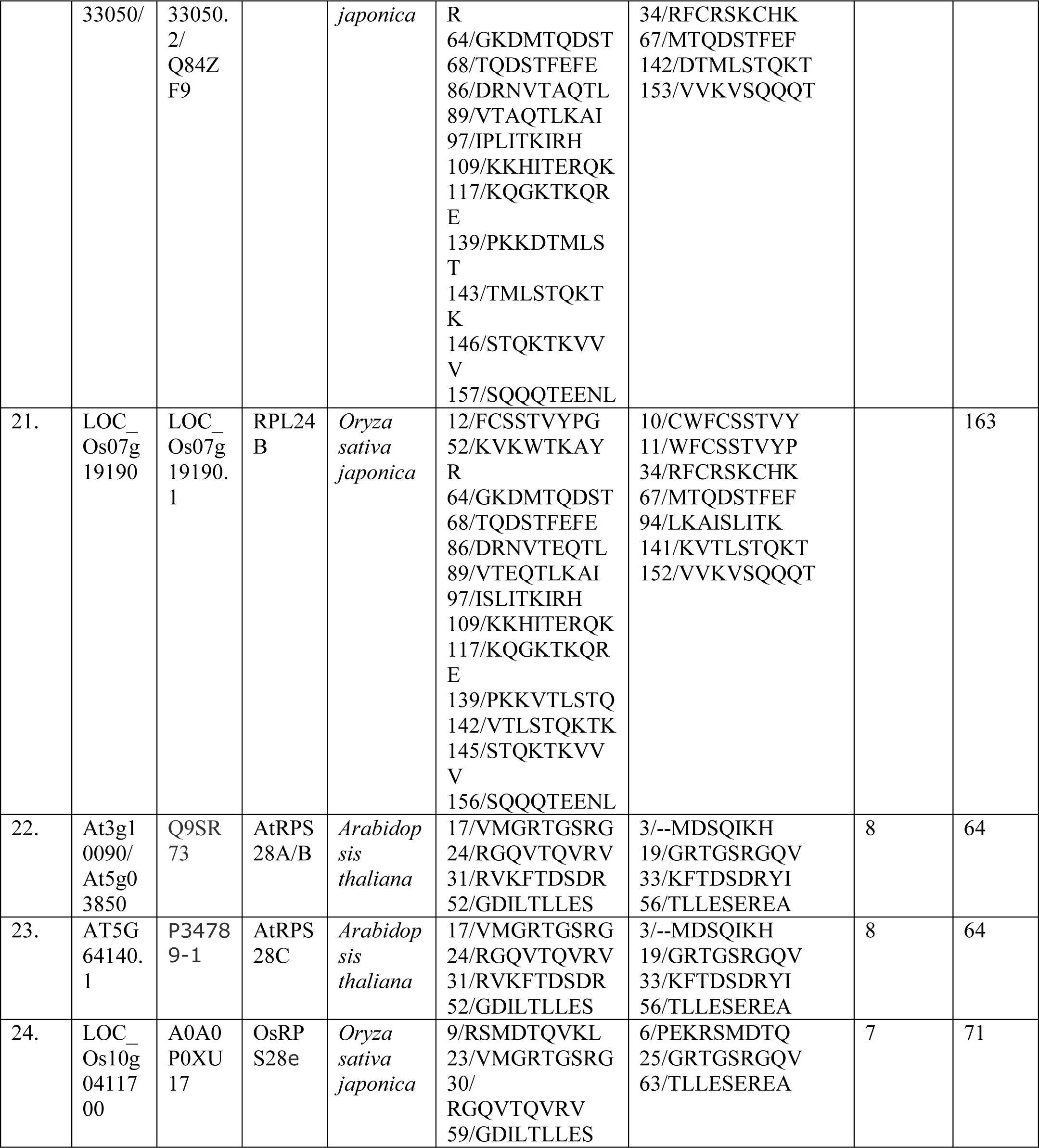
Identification of Ser/Thr phosphorylation sites in RPs mediated by AGC Kinases family (eg. PKC, PKG, RSK, PKA, PKB) The peptide sequences and Serine/ Threonine phosphorylation sites were identified in the Arabidopsis and Rice RPL6, RPL18, RPL23, RPL24 and RPS28 protein sequences and similaritywas made with the known peptide sequences and Serine/ Threonine phosphorylation sites of RPS6 protein using NetPhos 3.1 Server.

### 2.11. Expression studies of RP genes in the high *AtTOR* expressing rice lines

The expression analysis of genes encoding RPL and RPS was performed using qRT-PCR in two high expression lines TR-2.24 and TR-15.1 of rice in T_3_ generation (Bakshi et al. 2017). The rice transgenic lines were surface sterilized and allowed to grow on solid MS medium for 7d. The 7 DAG rice transgenic plants were used in the expression analysis. Total RNA was extracted and cDNA was synthesized as described earlier. The diluted cDNA was used for quantitative RT-PCR analysis of RPL and RPS genes.

### 2.12. Western blot analysis

Total protein was extracted from one month old plants of WT and T-DNA insertional Arabidopsis mutants of RP genes *rpl6* (CS16176), *rpl18* (SALK_134424), *rpl23A* (SALK_091329), *rpl24a* (SALK_064513), *rps28A* (SALK_094189), *Ats6k1* (SALK_113295.1) and *tor* (SALK_007846C) mutants using a standard phenol extraction method. The protein sample from the WT was taken as a positive control whereas protein samples from TOR and S6K1 mutants were taken as negative controls in the phosphorylation assay. The protein precipitate was dissolved in rehydration buffer (7 M Urea, 2 M Thiourea, 4% CHAPS and 30 mM DTT) and quantified by Bradford method with BSA standards. About 30 μg of total protein was loaded in SDS-PAGE and Western blot analysis. Human and Arabidopsis systems have a conserved phosphorylation site in p70 kDa ribosomal S6 Kinase 1 protein at Thr-389 residue (Xiong et al. 2013). We further performed S6K1 phosphorylation in Arabidopsis T-DNA insertional mutants for cytosolic ribosomal proteins genes *rpl6*, *rpl18*, *rpl23A*, *rpl24a*, *rps28A*, *s6k1* and *tor* to determine the cross-link between TOR and RPs using anti-human S6K1 antibody raised in mouse (anti phospho70kDa-S6K1-Thr(P)-389) (Cell Signaling Technologies, cat# 9206). The anti-70S6K1 (CST, cat# 2708S), anti-GAPDH (Santa Cruz, FL-335#SC25778) were used as loading controls. The membrane was incubated with secondary HRP-conjugated antibodies and signals were detected using a chemiluminescent method (ChemiDoc XRS, Bio-Rad).

## 3. Results

### 3.1. Molecular confirmation of *TOR*-OE transgenic plants

The seeds obtained from the primary *Agrobacterium*-treated plants were screened on PPT selection medium. The transgenic plants within 3-4 d after germination stayed green while non-transgenic and WT plants became bleached (Supplementary Fig. 1 & 2). After selection, plants were analyzed by PCR amplification of different elements present within the T-DNA using primers specific to the *bar* gene, CaMV35S promoter and *Arabidopsis thaliana* TOR gene (Supplementary Fig. 1c). The *Agrobacterium* transformation efficiency was calculated in T_1_ generation as described by Clough and Bent, (1998). Of about 20 plants that were infected with *Agrobacterium* carrying *AtTOR* binary vector, 13 were found to be positive on 10 mg l^−1^ PPT selection medium with a transformation efficiency of 65%. The *AtTOR* rice transgenic lines TR-2.24 and TR-15.1 were also similarly selected on 10 mg/L PPT selection medium with a transformation efficiency of 25.4% and were analyzed using PCR amplification (Bakshi et al., 2017).

### 3.2. Separation of *TOR*-OE lines

The T_3_-transgenic lines were germinated on PPT selection medium and based on their resistance and susceptibility on PPT, the ∑ Chi-Square (χ2) analysis was performed. The PPT resistant progenies exhibited Mendelian segregation ratio of 3:1 for the integrated T-DNA (Supplementary Table. 1). The T_3_ generation transgenic lines were separated on the basis of the level of *AtTOR* transcripts. The band intensity of *TOR* transcripts was observed on 1% agarose gel after Semi-Q PCR and the quantification was further confirmed by qRT-PCR (Fig. 1a, 1b, 1c & 1d). The transgenic lines ATR-1.4.12, ATR-ATR-1.4.27, ATR-1.7.10, ATR-1.2.23, ATR-1.6.21, ATR-3.5.42, ATR-3.8.41, and ATR-3.6.24 showed high band intensity. The lines, ATR-3.7.32, ATR-2.4.6 and ATR-2.4.28 exhibited bands with medium intensity and the lines, ATR-1.8.2 and ATR-4.1.11 had low intensity bands on the agarose gel.

The transcript levels of *TOR* were further analyzed by qRT-PCR. The shoot and root tissues of transgenic lines ATR-1.2.23 and ATR-1.7.10 exhibited ≥12-fold of *TOR* transcript level compared to WT. The transgenic lines, ATR-1.4.27 and ATR-1.4.12 exhibited more than five-fold up-regulation in shoots whereas the roots of these lines had more than 10-fold elevation. Similarly, transgenic lines ATR-3.6.24 and ATR-3.7.32 exhibited more than 5-fold and lines, ATR-2.4.6 and ATR-2.4.28 exhibited up to 4-fold upregulation in shoots and more than 5-fold in roots. On the basis of band intensity on agarose gel and transcript levels, the lines ATR-1.4.27 was considered as a high and ATR-3.7.32 was considered as a medium expression line. The selected two lines ATR-1.4.27 and ATR-3.7.32 were used for further investigations. The *AtTOR* rice transgenic lines TR-2.24 and TR-15.1 were considered as high expression lines having transcript levels up to 30-fold in shoots and up to 80-fold in roots (Bakshi et al., 2017).

### 3.3. Response of *TOR*-OE lines to various stress treatments

To address the roles of TOR in response to various abiotic stresses the 15 d old plants representing two *TOR*-OE Arabidopsis transgenic lines ATR-1.4.27 and ATR-3.7.32 were subjected to osmotic and salt stress treatments. In our previous report (Bakshi et al., 2017), we observed that the *TOR* gene functions in a dose dependent manner where the level of *TOR* gene expression was directly linked to the plant phenotypes and increased tolerance to abiotic stress treatments. Therefore, we selected high *TOR* expression lines of Arabidopsis and rice in our present study. The *TOR*-OE transgenic plants along with the WT were subjected to NaCl (150 mM), PEG (7%), sorbitol (200 mM) and mannitol (100 mM) stress treatments (Fig. 2a, 2b, 2c & 2d). After 7 DAG on stress treatments, the fresh weight of 10 plants and primary root length (using a 1 cm scale bar) were recorded in three biological replicates (Fig. 3& 4).

**Figure 3.**
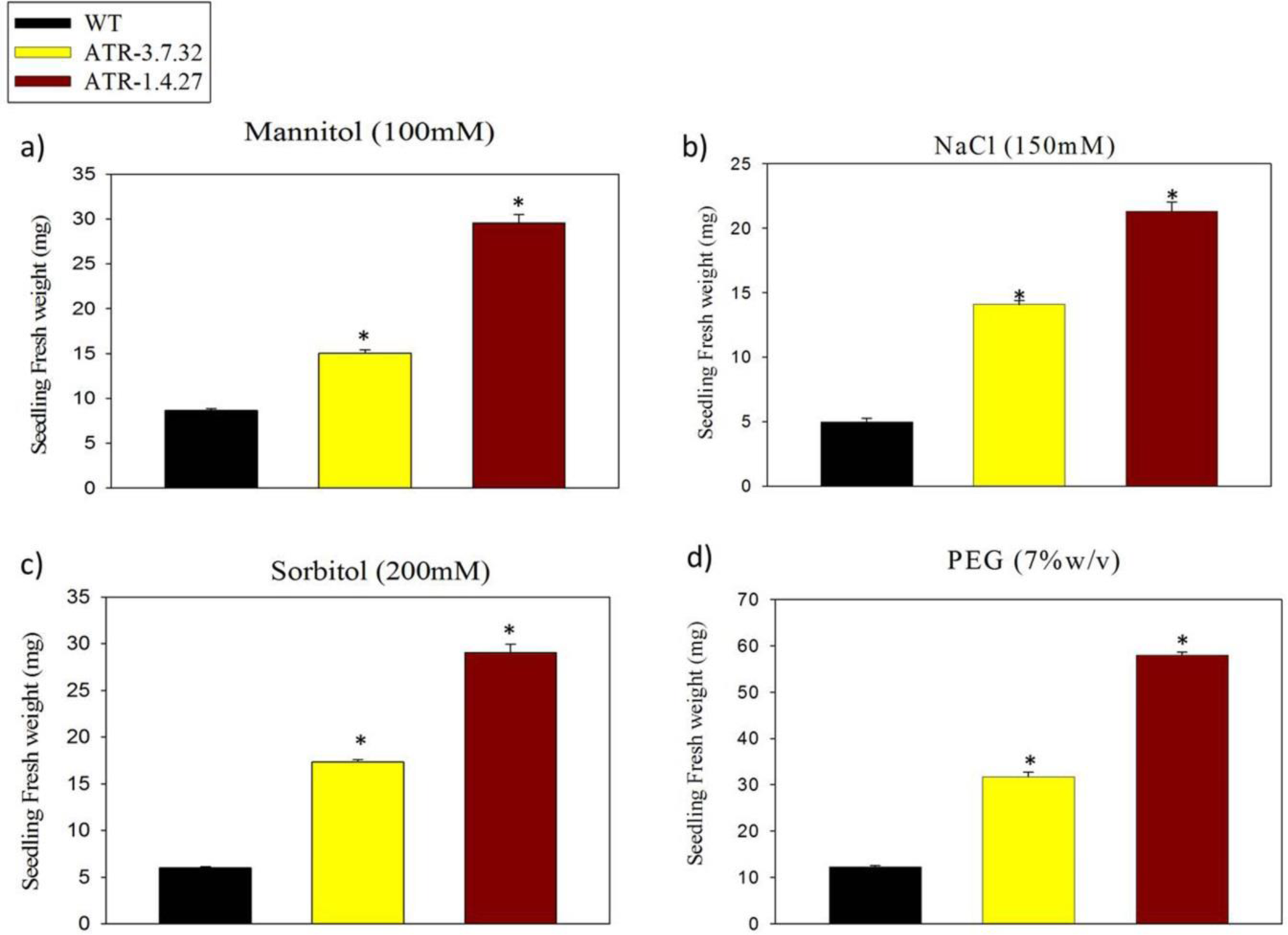
Seedling fresh weight (mg) of *TOR*-OE Arabidopsis lines in abiotic stress treatments. The fresh weight of ten plants of transgenic *TOR-*OE (ATR-1.4.27 and ATR-3.7.32) and WT were recorded in three replicates after 7 days of every treatment. (a) After mannitol treatment the two transgenic lines had increased shoot growth compared with the treated WT. (b) The shoots of WT plants became completely bleached at high salt (NaCl 150mM) concentration after 7 DAG treatment, whereas the two transgenic lines exhibited healthy growth with high shoot and root growth. (c) Sorbitol (200 mM) and (d) PEG treated transgenic plants also exhibited increased shoot and root growth, whereas the growth of WT plants was retarded under high osmotic stress. The two lines (ATR-1.4.27 and ATR-3.7.32) had increased shoot biomass compared with the WT seedling.

Although, the phenotypic characterization of *TOR* overexpression was previously reported in *Arabidopsis thaliana* (Deprost et al. 2007; Ren et al. 2011), their performance and phenotypic variation under high osmotic and salt stress have not been evaluated so far. In the present investigation, we have performed treatments on *TOR*-OE Arabidopsis seedlings with various chemicals that induce stress (Sorbitol, Mannitol, PEG and NaCl). The phenotypes of Arabidopsis *TOR*-OE lines showed by Ren et al (2011) had defective root and shoot growth at a very high level of *TOR* expression, ∼80 fold relative to the wild-type control. Also, the phenotypic aberrations observed by Ren et al. (2011) were at the stage of maturity in Arabidopsis *TOR* overexpression lines. To avoid these phenotypic abnormalities we selected the early stage (7 DAG) *TOR*-OE plants with highest *TOR* expression of 15 folds for osmotic and salt stress treatments. The high expression line, ATR-1.4.27 (>10-fold in shoots) and medium expression line, ATR-3.7.32 (∼7-fold in shoots) exhibited significantly increased shoot weight and root length compared to the WT in response to all the treatments. The fresh weight of ten plants of high expression line (30 mg) was also increased in response to mannitol treatment compared to the medium line and WT (Fig. 3a). The mannitol treated ATR-1.4.27 line exhibited increased root length (5.5 cm) whereas the mediumexpression line ATR-3.7.32 and WT showed no significant change in root length (≥2 cm; Fig. 4a). The NaCl treatment caused bleaching of WT plants with retarded growth whereas the other two lines exhibited healthy shoot and root growth. Both the transgenic lines ATR-1.4.27 and ATR-3.7.32 showed increased root length up to 3.0 cm and the fresh weight was more than 20 mg per ten plants under NaCl treatment compared with the WT, which had a root length of 1.2 cm and 4.5 mg of fresh weight (Fig. 3b & 4b). Similarly, the Sorbitol and PEG treatments in the two transgenic lines also resulted in increased root length and fresh weight compared with the WT (Fig. 3c, 3d, 4c & 4d). Under sorbitol and PEG treatments, the transgenic lines also showed increased lateral root formation, whereas the WT exhibited root growth inhibition. The data obtained from phenotypic analyses were plotted using the mean of three biological replicates in response to each treatment.

**Figure 4.**
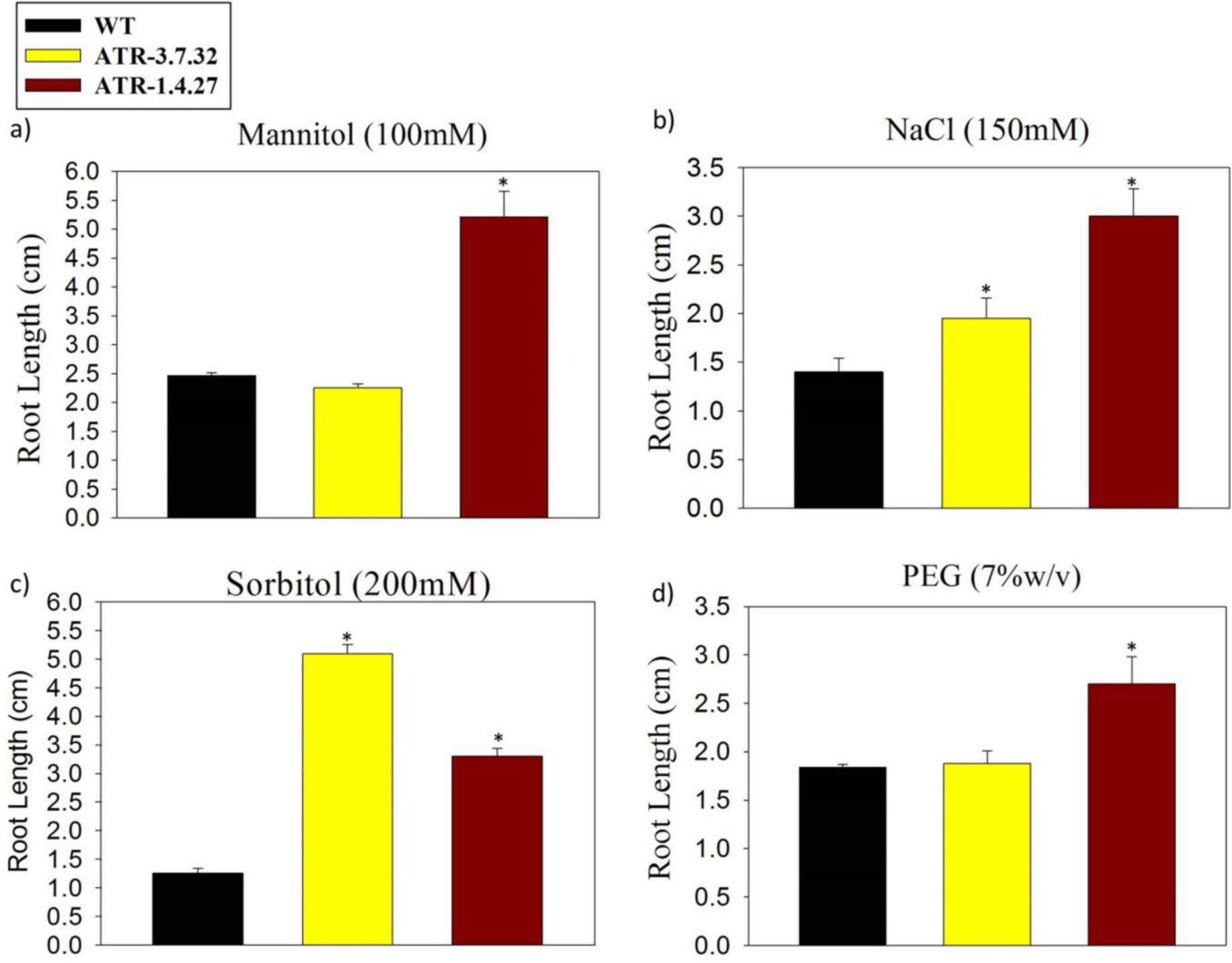
Root length measurement of *TOR*-OE Arabidopsis lines. The root length was measured in three biological replicates using 1 cm scale bar after 7 DAG of treatment. (a) The high expression line (ATR-1.4.27, 12-fold) showed increased root length in response to mannitol treatment and the medium line exhibited root length similar to the WT. (b) Both the transgenic lines (ATR-1.4.27 and ATR-3.7.32) also exhibited increased root lengths in NaCl treatment compared with the WT. (c) The transgenic lines (ATR-1.4.27 and ATR-3.7.32) showed increased primary and lateral root growth under sorbitol treatment compared with the WT. (d) Treatment of high and medium lines, ATR-1.4.27 and ATR-3.7.32 with PEG (7% w/v) resulted in reduced root growth in WT plants, in contrast the *TOR*-OE lines ATR-1.4.27 (3 cm) exhibited increased primary and lateral root growth The line ATR-3.7.32 and WT had root lengths of (≤ 2 cm) but the line, ATR-3.7.32 had increased lateral root growth.

### 3.4. Estimation of chlorophyll content and percent degradation

Total chlorophyll content is likely to be affected by the stress environment, which is also directly related to the photosynthestic efficiency and plant performance under abiotic stress conditions (Bouvier et al., 2005). The chlorophyll tends to degrade under oxidative stress and measurement of cholorophyll content is an important physiological parameter to demonstrate stress tolerance in a plant (Hasegawa et al., 2000). We observed that the *TOR* overexpressing Arabidopsis plants had high chlorophyll content and low percentage of chlorophyll degradation when treated with NaCl (150 mM), PEG-8000 (7%), sorbitol (200 mM) and Mannitol (100 mM). The two transgenic lines, ATR-1.4.27 and ATR-3.7.32 had increased chlorophyll-a content up to 25 µg mg^−1^ and 20 µg mg^−1^, respectively in untreated conditions compared with the WT Arabidopsis plants, which had a concentration of 17 µg mg^−1^. The Chl-a content in two lines (ATR-1.4.27 and ATR-3.7.32) was 10 µg mg^−1^ and 14 µg mg^−1^, respectively under NaCl treatment compared with the WT, which exhibited only 2 µg mg^−1^. The sorbitol treated transgenic plants had chlorophyll-a content ranging from 20-25 µg mg^−1^, whereas the WT had 17 µg mg^−1^. The mannitol treatment also had reduced chlorophyll-a content in WT (up to 5 µg mg^−1^) compared with the untreated conditions. The two transgenic lines had 15 µg mg^−1^ to 20 µg mg^−1^ of Chl-a (Fig. 5a). The Chl-a content was also high in the two transgenic lines in response to 7% PEG. The high expression transgenic line, ATR-1.4.27 had more than 15 µg mg^−1^ and the medium expression line, ATR-3.7.32 had below 15 µg mg^−1^ of Chl-a content compared with the PEG-treated WT which had 11 µg mg^−1^.

**Figure 5.**
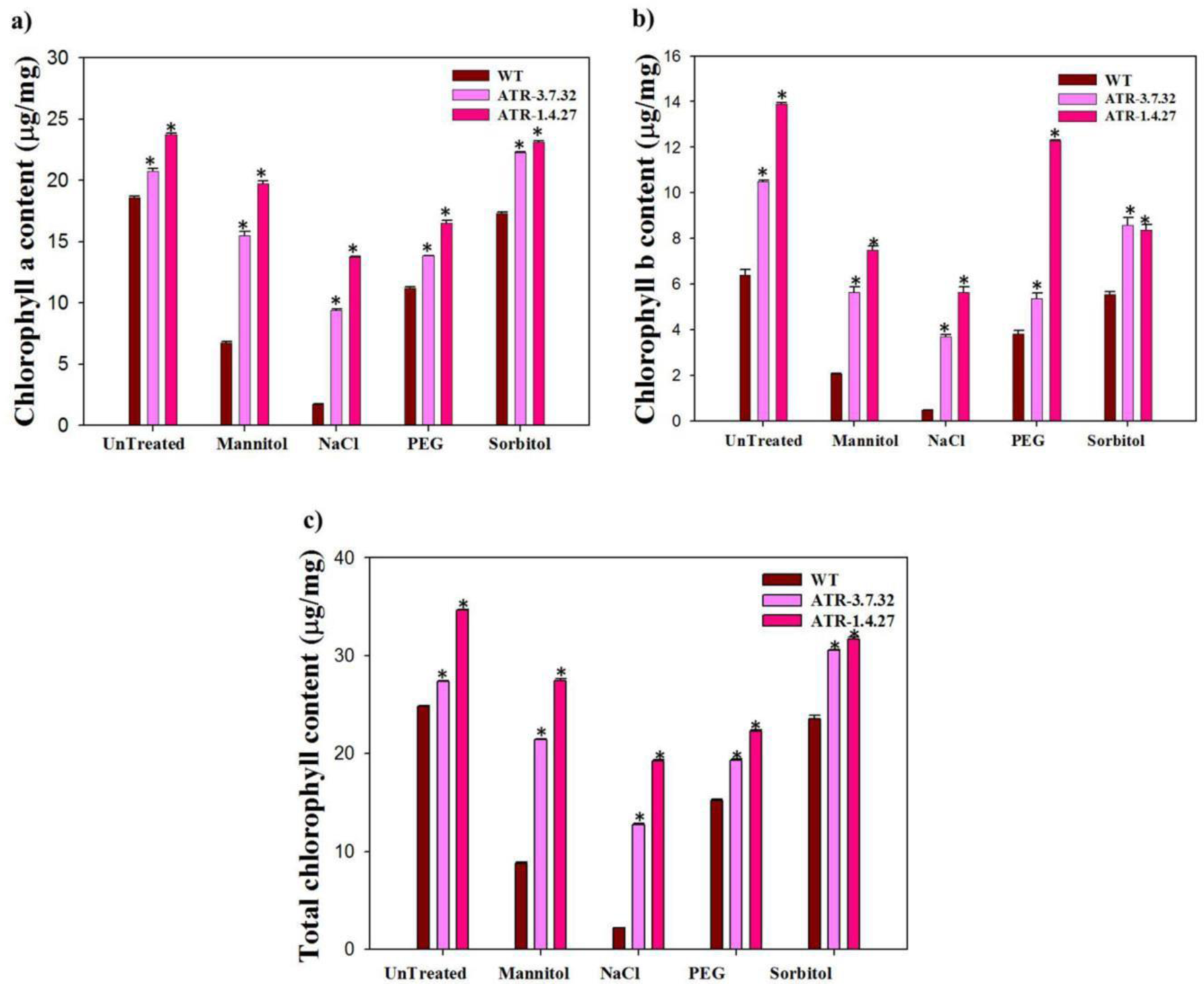
Estimation of Chlorophyll a, b and total contents. (a) Chlorophyll-a, (b) Chlorophyll-b, and (c) total chlorophyll contents were measured under both treated and untreated plants of high (ATR-1.4.27), medium (ATR-3.7.32) lines and WT. The mean values of chlorophyll data with ±SE was represented with asterisks (*) and were considered statistically significant at *P*<0.05.

Similarly, the Chl-b content in untreated medium expression line (ATR-3.7.32) was also as high as 11.5 µg mg^−1^ and the high expression line, ATR-1.4.27 had 14 µg mg^−1^, whereas the WT had 6.5 µg mg^−1^. In response to NaCl, the Chl-b was also as high as 6 µg/ml in ATR-1.4.27 and 4 µg mg^−1^ in ATR-3.7.32, whereas the WT had <1 µg mg^−1^ under NaCl treatment. Also, the Chl-b content was high in two transgenic lines under PEG, sorbitol and mannitol treatments ranging from 4 µg/ml to 12 µg mg^−1^ compared with WT ranging from 2 µg mg^−1^ to 5 µg mg^−1^ (Fig. 5b).

The chlorophyll degradation percentage was calculated with the values of chlorophyll contents estimated before and after the abiotic stress treatments. The NaCl treatment had resulted in more than 90% degradation of Chl-a, b and total contents in the WT plants, whereas the two transgenic lines had degradation ranging from 40 to 50%. The Chl content was degraded up to 60% in WT under mannitol treatment and the transgenic lines had a degradation of 20% in Chl-a and total contents, whereas the Chl-b degradation was more than 40% (Fig. 6a, 6b, 6c & 6d).

**Figure 6.**
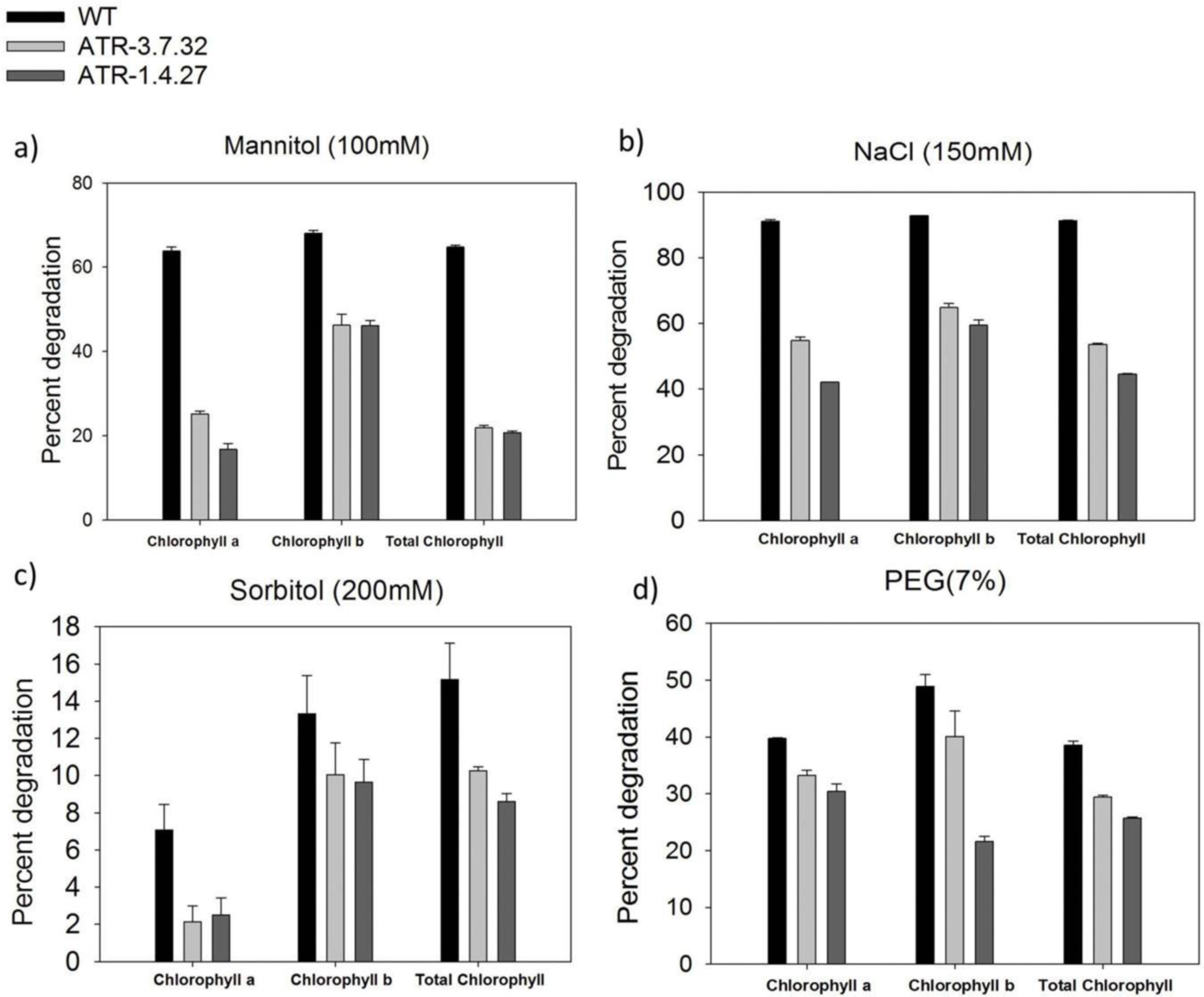
Chlorophyll degradation percentage in *TOR*-OE Arabidopsis plants. (a) Percent degradation of Chl-a, b and total contents after NaCl stress treatment was low in two selected lines compared with the WT. (b) **∼**20% of percent degradation of Chl-a and total chlorophyll content was observed in two transgenic lines after mannitol treatment, whereas the WT had >50% degradation of Chl-a, b and total chlorophyll contents (c and d) similarly, under PEG and sorbitol treatments, the high expression lines had low percent degradation of three extracts.

### 3.5. Proline estimation

Plants accumulate osmolytes such as proline, proline betaine, glycine betaine, mannitol and sorbitol to reduce cell damage caused by the reactive oxygen species (ROS) produced under osmotic and salinity stress conditions (Hare and Cress 1997). Proline is a low molecular weight osmolyte, which is accumulated in response to osmotic and salinity stresses in plants (Hasegawa et al., 2000). The proline content was estimated in two *TOR*-OE lines ATR-3.7.32 and ATR-1.4.27 in comparison with the WT plants after all the stress treatments. The untreated ATR-3.7.32 and ATR-1.4.27 plants had proline contents of 4.5 µg mg^−1^ and 5 µg mg^−1^, respectively and the WT seedling had 3.8 µg mg^−1^ proline. The NaCl treatment in lines ATR-1.4.27 and ATR-3.7.32 had increased proline accumulation up to 16 µg mg^−1^ and 14 µg mg^−1^, respectively when compared with the WT which had 7 µg mg^−1^ proline. The PEG and Mannitol treated transgenic plants accumulated more than 30 µg mg^−1^ proline after treatment compared to the WT (10 µg mg^−1^ proline). The sorbitol treatment had also increased the accumulation of proline up to 14 µg mg^−1^ in transgenic lines compared with the WT which had 5 µg mg^−1^ proline (Fig. 7).

**Figure 7.**
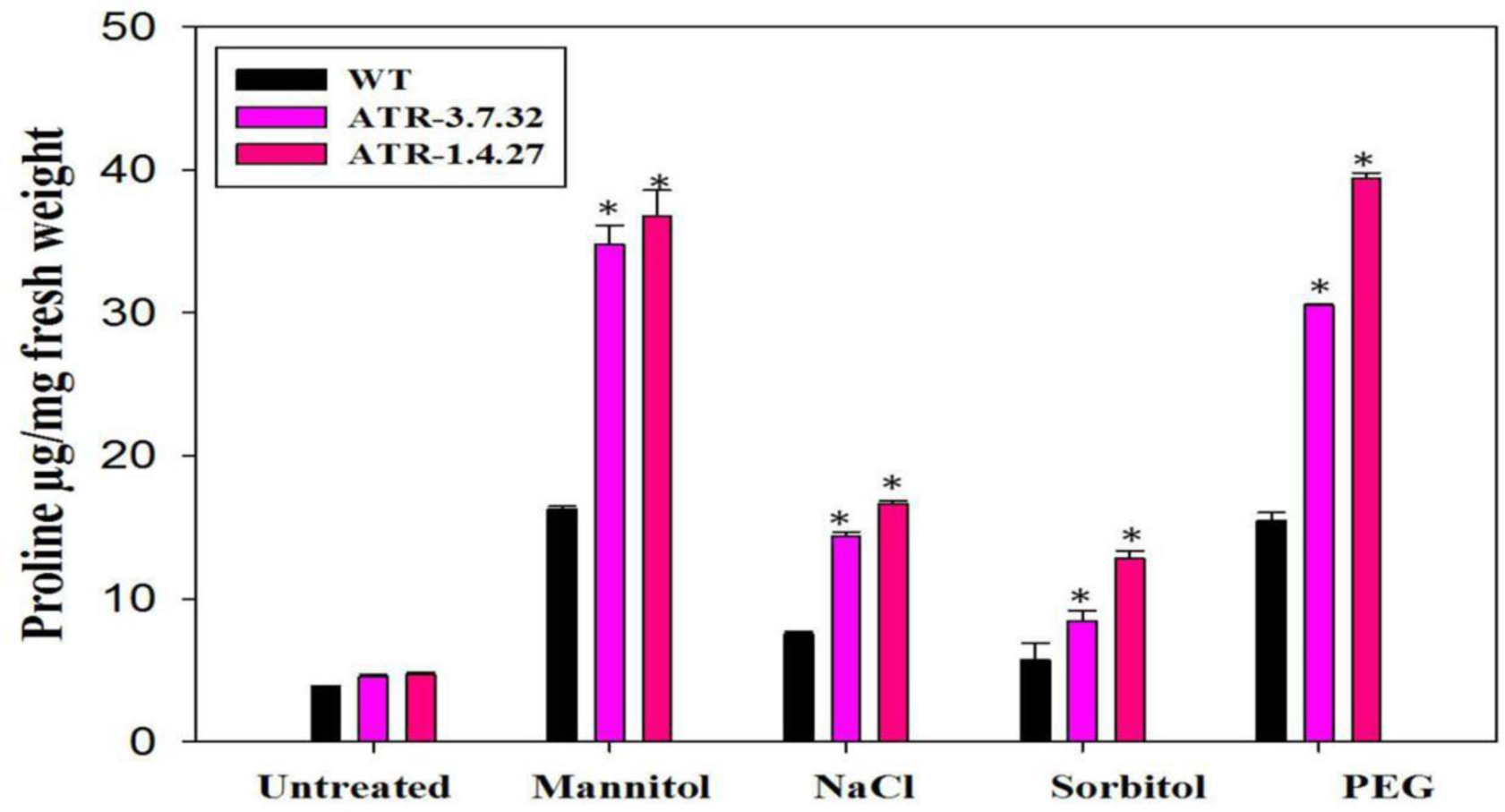
Estimation of Proline content in *TOR*-OE Arabidopsis plants. The proline content was estimated before and after stress treatments in *TOR*-OE transgenic lines and WT. Representation of proline (a) in untreated and (b) NaCl-treated (150 mM) transgenic lines and WT. (c, d, and e) Proline content in PEG (7% w/v), mannitol (100 mM) and sorbitol (200 mM) treated lines (ATR-1.4.27, ATR-3.7.32) and WT. The proline content was estimated in three biological replicates and the mean was plotted with ± SE. The statistical significance was calculated based on one-way ANOVA at *P* value <0.05 and represented with asterisks in the graphs.

### 3.6. Expression of stress-inducible genes

The expression patterns of eight stress-inducible genes, *AtERD*11 (Early response to dehydration 11), *AtSAMDC* (S-Adenosyl methionine decarboxylic carboxylase), *AtAPX*1 (Ascorbate peroxidase), *AtERF*5 (Ethylene responsive Factor), *AtCSD*1 (Cu/Zn super oxide dismutase), *AtMSD*1 (Mn superoxide dismutase), *AtSOS*1 (Salt overly sensitive 1) and *AtCATALASE* were analyzed in treated and untreated plants of *TOR*-OE lines, ATR-3.7.32 and ATR-1.4.27 and WT (Fig. 8). The overexpression of *TOR* was associated with the upregulation of all the analyzed stress-inducible genes. The *AtERD11* gene, which encodes a glutathione S-transferase in Arabidopsis, is responsive to dehydration stress (Kiyosue et al. 1993). The transcripts of *AtERD*11 in transgenic lines (ATR-1.4.27 and ATR 3.7.32) were upregulated more than 1.5-fold in NaCl and sorbitol treatments, whereas the PEG-treated plants had the transcript level of *AtERD*11 up to ∼15-fold. Also, the mannitol treatment had increased the expression of *AtERD*11 up to 5-fold in these two lines (Fig. 8a). The ROS-mediated TOR signaling has been reported as a switch for regulation of root growth and autophagy (Yokawa et al. 2015). The At*SOS*1, which regulates Na+ / H+ homeostasis under stress conditions was also highly up-regulated in the two lines under all the treatments, whereas the *AtSOS*1 transcript was increased by more than 10-fold under NaCl treatment in line, ATR-1.4.27 (Fig. 8b). The genes, *AtCATALASE*, *AtAPX*1, *AtMSD*1 and *AtCSD*1, which are involved in the regulation of ROS levels in the cell were also highly up-regulated in two selected lines (Fig. 8c, 8d, 8g & 8h). The transcript level of *AtCATALASE* was increased up to 7-fold under NaCl, >1.5-fold in response to sorbitol and PEG and up to 6-fold under mannitol treatments, respectively (Fig. 8c). Both the lines exhibited significant upregulation of *AtSAMDC* & *AtERF5* genes with more than two fold in all the stress treatments, which however, showed more elevation in response to PEG treatment (Fig. 8e & 8f). The *AtAPX*1, a ROS scavenger catalyzes the conversion of ascorbate and hydrogen peroxide to less toxic mono-dehydroascorbate and H_2_O. The expression of *AtAPX*1 was up-regulated to more than 15-fold under sorbitol, mannitol and PEG treatments (Fig. 8g). The *AtMSD1* and *AtCSD1* expression was also enhanced in the two transgenic lines in all the treatments compared with the treated WT (Fig. 8d & 8h). The expression of *AtMSD*1 was also up-regulated more than 2-fold under sorbitol and mannitol treatment and more than 6-fold under NaCl and PEG treatments. Similarly, the sorbitol and mannitol treated transgenic plants had an elevation of *AtCSD*1 transcripts up to 10-fold suggesting that there was ROS regulation that is in line with the overexpression of *TOR*.

**Figure 8.**
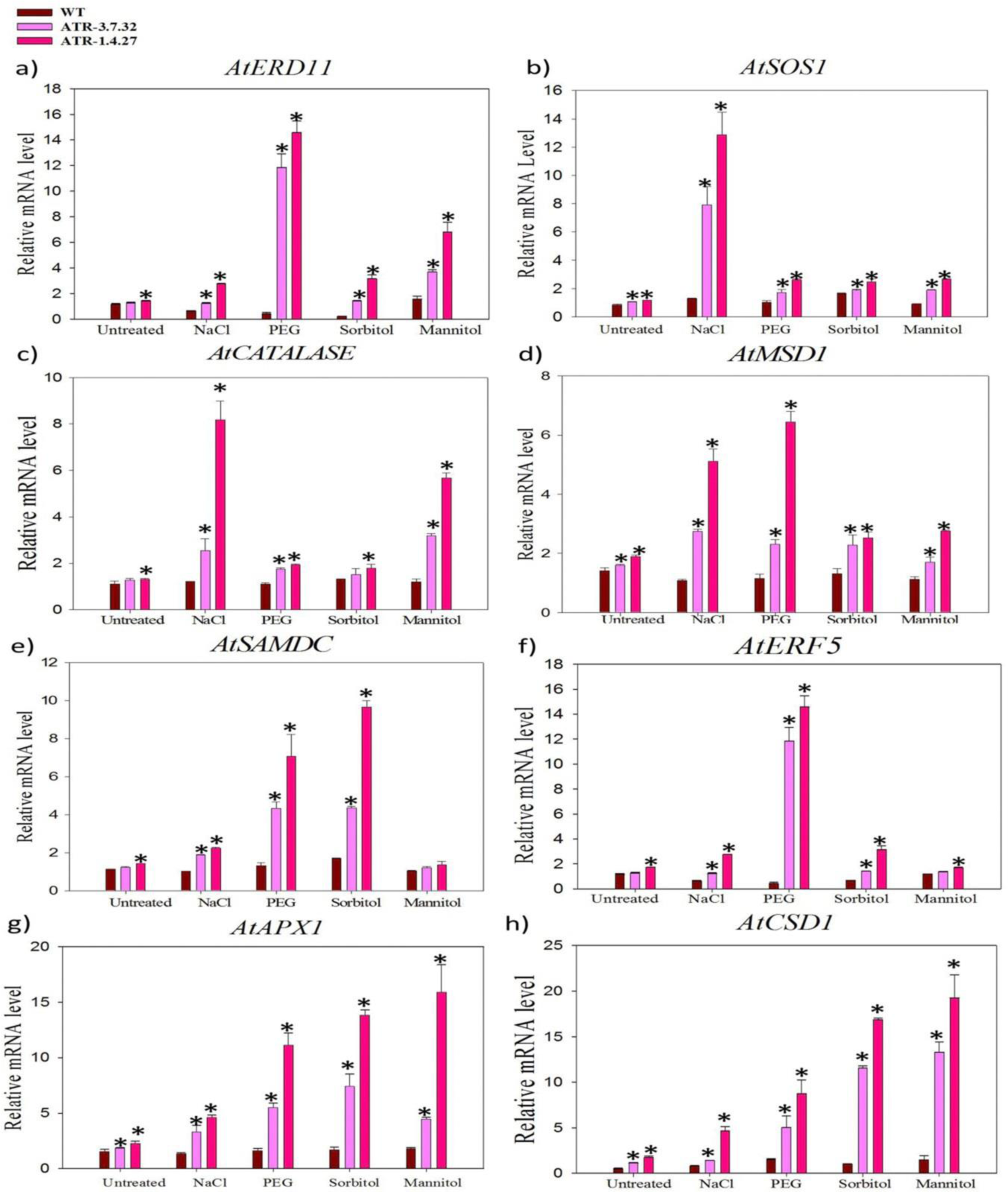
Transcriptional regulation of stress-specific genes. The expression of stress-responsive genes was analyzed in treated and untreated plants of ATR-1.4.27 and ATR-3.7.32 lines in comparison with the WT. The expression of stress-specific genes such as (a) *AtERD11* (b) *AtSOS1* and (c) *AtCATALASE* was studied in root and shoots tissues. The stress-specific genes such as (d) *AtMSD1* (e) *AtSAMDC* and (f) *AtERF5* were studied in roots and shoots of two selected lines. The up-regulation of these genes in all the stress treatments indicates the role of TOR in transcriptional regulation of stress-specific genes. The abiotic treated and untreated plants of the high and medium TOR expression lines also exhibited up-regulation of stress-inducible genes, (g) *AtAPX1*, (h) *AtCSD1* in roots and shoots. The mean of three biological replicates was used in qRT-PCR data and Arabidopsis specific Actin2 was used as an endogenous reference gene for normalization. The relative expression was considered statistically significant at P value <0.05 (represented with asterisks ‘*’) based on one-way ANOVA in all the analyzed genes.

The 7 DAG plants of the two high *AtTOR* expression rice lines TR-2.24 and TR-15.1 (Bakshi et al., 2017) were also similarly treated with PEG and NaCl and the transcript levels of stress-responsive genes were analyzed in root and shoot tissues using qRT-PCR. We observed that the genes involved in osmotic protection such as *OsNADPH*1, *OsALDH*2a, and OsLEA3-1 were highly upregulated in the high expression lines, TR-2.24 and TR-15.10 in both treated and untreated conditions (Bakshi et al., 2017). We also observed that the genes involved in the ROS sequestration such as Alternative oxidase (*OsAOX*1a) in rice lines TR-2.24 and TR-15.1 and Ascorbate peroxidase (*AtAPX*1) in the Arabidopsis lines ATR-1.4.27 and ATR-3.7.32 were significantly upregulated in the PEG and NaCl treatment. The enhanced expression of genes related to ROS regulation along with increased root phenotype observed after stress treatments in the Arabidopsis and rice transgenic plants suggest a possible link between ROS-*TOR* signaling in plants.

### 3.7. Expression analysis of Ribosomal Protein Large (RPL) and Small subunit (RPS) genes in rice

Each ribosomal protein is encoded by its corresponding orthologous gene. RPs are heterogeneous in nature i.e. each RP is encoded by one of the multiple homologous partners for a given protein (Wool et al. 1979), which is determined by the tissue and the environmental conditions of the plant. The TOR-mediated regulation of rRNA synthesis, ribosome biogenesis and protein synthesis has been reported in Arabidopsis (Ren et al. 2011), but the regulation of genes encoding RPs by TOR has not been analyzed in plants. Our comparative transcript analysis of genes encoding large and small ribosomal subunit proteins in previously generated high *AtTOR* expressing rice transgenic lines, TR-2.24 and TR-15.1 clearly suggested that TOR modulates the transcription of RP genes (Bakshi et al., 2017)..

We have previously observed that the ectopic expression of *AtTOR* had increased water-use efficiency and yield related attributes in rice (Bakshi et al. 2017). Previously, the effect of TOR inactivation on the expression of Arabidopsis ribosomal proteins has been investigated by using transcriptomic and phosphoproteomic analysis (Dobrenel et al. 2016). The RP gene expression data of AtTOR expressing rice lines TR-2.24 and TR-15.1 obtained by qRT-PCR analysis was compared with the available transcriptomic reports of RP genes by other groups in TOR-RNAi lines of Arabidopsis (Supplementary Fig. 3a & 3b; Dobrenel et al., 2016). The inactivation of TOR in two TOR-RNAi lines of Arabidopsis resulted in coordinated decrease of the transcription and translation of plastidic ribosomal protein genes, whereas the genes coding for cytosolic ribosomal proteins were interestingly upregulated (Dobrenel et al. 2016). The chloroplast degradation occurred during autophagy induced by the TOR inactivation may also cause down-regulation of plastidic ribosomal genes. Simultaneously, the fact that cytosolic ribosomal protein genes are induced under abiotic and biotic stress conditions, which is also a consequence of the inhibition of TOR. However, the DEG data observed by Dong et al., (2015) in the asTORi Arabidopsis lines showed downregulation of 114 RP genes and 1 up-regulated gene associated with ribosome pathway. Doberenel et al., (2016) observed that the transcript levels of the cytoplasmic genes *RPS2, RPS3 RPS 6, RPS9, RPS10, RPS11, RPS12, RPS13, RPS15, RPS16, RPS17, RPS18, RPS19, RPS21, RPS26, RPS27, RPS27a, RPS28, RPL3, RPL 4, RPL 5, RPL 6, RPL 7a, RPL 8, RPL 9, RPL 10. RPL 10a, RPL 11, RPL 12, RPL 14, RPL 15, RPL 18, RPL 18a, RPL 19, RPL 21, RPL 22, RPL 23, RPL 23a, RPL 24, RPL 27, RPL 27a, RPL 28, RPL 30, RPL 31, RPL 32, RPL 34, RPL 35, RPL 35a, RPL 36, RPL 36a, RPL 37, RPL 37a, RPL 38, RPL 39, RPL 40* and *RPL41* were significantly upregulated in TOR RNAi lines, whereas the expression of genes *RPS5, RPS 8, RPS 14, RPS 20, RPS 23, RPS 29, RPS 30, RPL7, RPL13, RPL13a, RPL17, RPL26* and *RPL29* were either slightly modulated or remained unchanged (Dobrenel et al., 2016).

Although th involvement of TOR signaling in modulation of RP genes transcription has been demonstrated earlier in Arabidopsis (Kojima et al. 2007; Ren et al. 2011; Dong et al. 2015; Dobrenel et al. 2016), no such reports areavailable in crop plants. Further, based on the previous studies on the extra-ribosomal function of RPs in response to abiotic stresses in rice (Moin et al. 2016; Moin et al. 2017; Saha et al. 2017; Bakshi et al. 2019) and to understand the effect of constitutive expression of *AtTOR* on regulation of RPs transcription, we performed expression analysis of RPL and RPS genes in two high *AtTOR* expressing rice lines, TR-2.24 and TR-15.1 (Bakshi et al. 2017). The expression analysis of all the rice RP genes in two high expression lines (TR-2.24 and TR-15.1) suggested a TOR-mediated regulation of RP genes in plants. The transcript levels of almost all the RP genes were elevated in the rice transgenic lines. The genes, *RPL4*, *RPL14*, *RPL18A*, *RPL19.3*, *RPL36.2*, *RPL51*, *RPS3A*, *RPS6*, *RPS6A*, *RPS25A* and *RPS30* became highly up-regulated in the rice transgenic plants that express *AtTOR* ectopically (Fig. 9a & 9b). The transcript levels of *RPL18A*, *RPL19.3*, *RPL51*, *RPS25A* and *RPS30* were significantly increased up to ten fold and *RPL4*, *RPL14*, *RPL24B*, *RPL26.1*, *RPL30e*, *RPL38A*, *RPL44*, *RPS3A*, *RPS6*, *RPS6A*, *RPS27* and *RPS27a* were upregulated more than five-fold. The *RPL6*, *RPmL6* (mitochondrial), *RPnL6* (nucleolar), *RPL24*, *RPL18*, *RPL23A* and *RPS28A* expression was also enhanced up to two-fold in TR-2.24 and TR-15.1 rice transgenic lines. The genes *RPL5, RPL15, RPL23a, RPL24, RPL27.3, RPL37, RPS18a, RPS18b, RPS23a* and *RPS24* were slightly upregulated less than 2 fold. Altogether, these data strongly suggest a positive correlation between the TOR and RP genes transcription and translation in plants.

**Figure 9.**
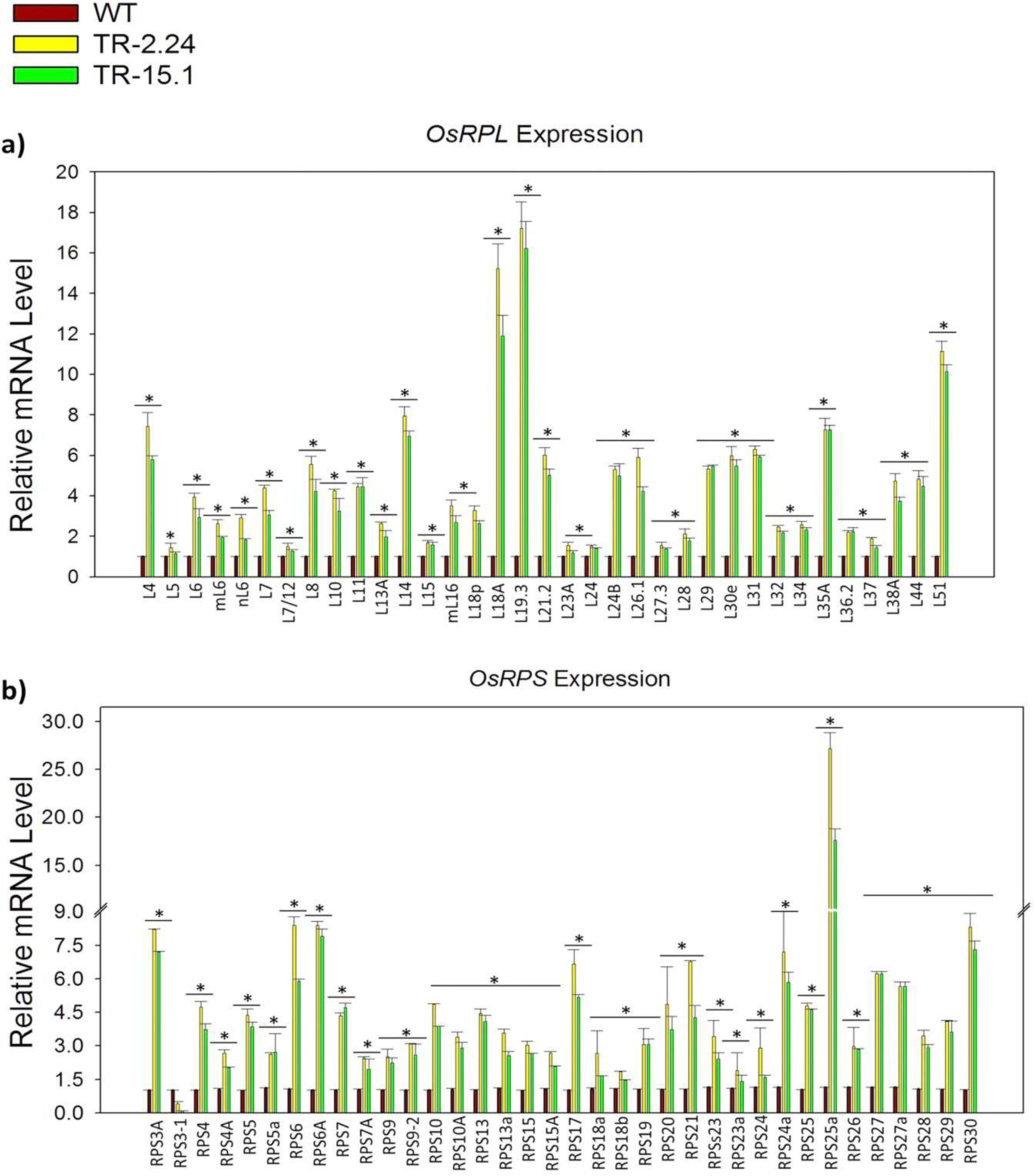
Transcriptional regulation of Ribosomal Protein Large subunit (RPL) and Small subunit (RPS) genes. The 7 DAG plants of two high *AtTOR* expressing rice transgenic lines TR-2.24 and TR-15.1 were used to analyze the expression of Ribosomal Protein Large subunit (RPL) and Ribosomal Protein Small subunit (RPS) genes. The WT plants were used as control. (a) Expression analysis of rice RPL genes in two high AtTOR expression lines of rice, TR-2.24 and TR-15.1. The *RPL4*, *RPL14*, *RPL18A*, *RPL19.3*, *RPL36.2*, *RPL51* genes were highly upregulated upto 20-fold in both the transgenic lines. (b) Expression analysis of rice RPS genes in two transgenic lines. The significant upregulation of RPS genes transcript in two transgenic lines was observed, where the *RPS3A*, *RPS6*, *RPS6A*, *RPS25A* and *RPS30* genes were highly upregulated more than 7-fold in transgenic plants. The fold change was normalized using ΔΔC_T_ method relative to the WT plants. Rice *Actin* (*Act*1) was used as an internal control. Three biological and three technical replicates were included in this study.

### 3.8. Ribosomal protein inhibition modulates feedback regulation of S6K1 phosphorylation

To gain more insights into the involvement RPs in phosphorylation of S6K protein, we performed a phosphoproteomic analysis of S6K1 in Arabidopsis insertional mutants for some of the important ribosomal proteins and observed that the mutation of RPL and RPS in Arabidopsis resulted in differential regulation of S6K1 phosphorylation in Arabidopsis. Simultaneously, the mutation in RPs also resulted in the loss of total S6K1 stability along with its phosphorylation. The TOR and S6K1 mutants were used as negative controls and both mutants showed inhibited S6K1 phosphorylation and reduced stability of total S6K1 protein. However, the TOR mutant showed slight phosphorylation of S6K1 protein in comparison with the S6K1 mutant, in which the phosphorylation was completely inhibited. The *rpl6* mutants had equally phosphorylated S6K1 protein, whereas the phosphorylation of S6K1 in *rpl23a*, *rpl24*, *rpl24a* and *rps28a* was slightly reduced. The stability of total S6K1 protein in *rpl6*, *rpl18a* and *rpl23a* was almost similar as in the WT sample, whereas the *rpl6*, *rpl18a* and *rpl23a* mutants had increased stability of total S6K1 protein. The mutation of *rpl18* had completely inhibited S6K1 phosphorylation and the *rpl24a* mutant had moderate inhibition, whereas the mutation in *rpl23* and *rps28* had no effect on S6K1 phosphorylation (Fig. 10a & 10b). The WT Col0 protein was used as a positive control for phosphorylation study and GAPDH was used as an endogenous loading control (Fig. 10c). Our phosphoproteomic results clearly suggested that ribosomal proteins are interlinked and involved in regulation of S6K1 phosphorylation in plants as in the animal systems.

**Figure 10.**
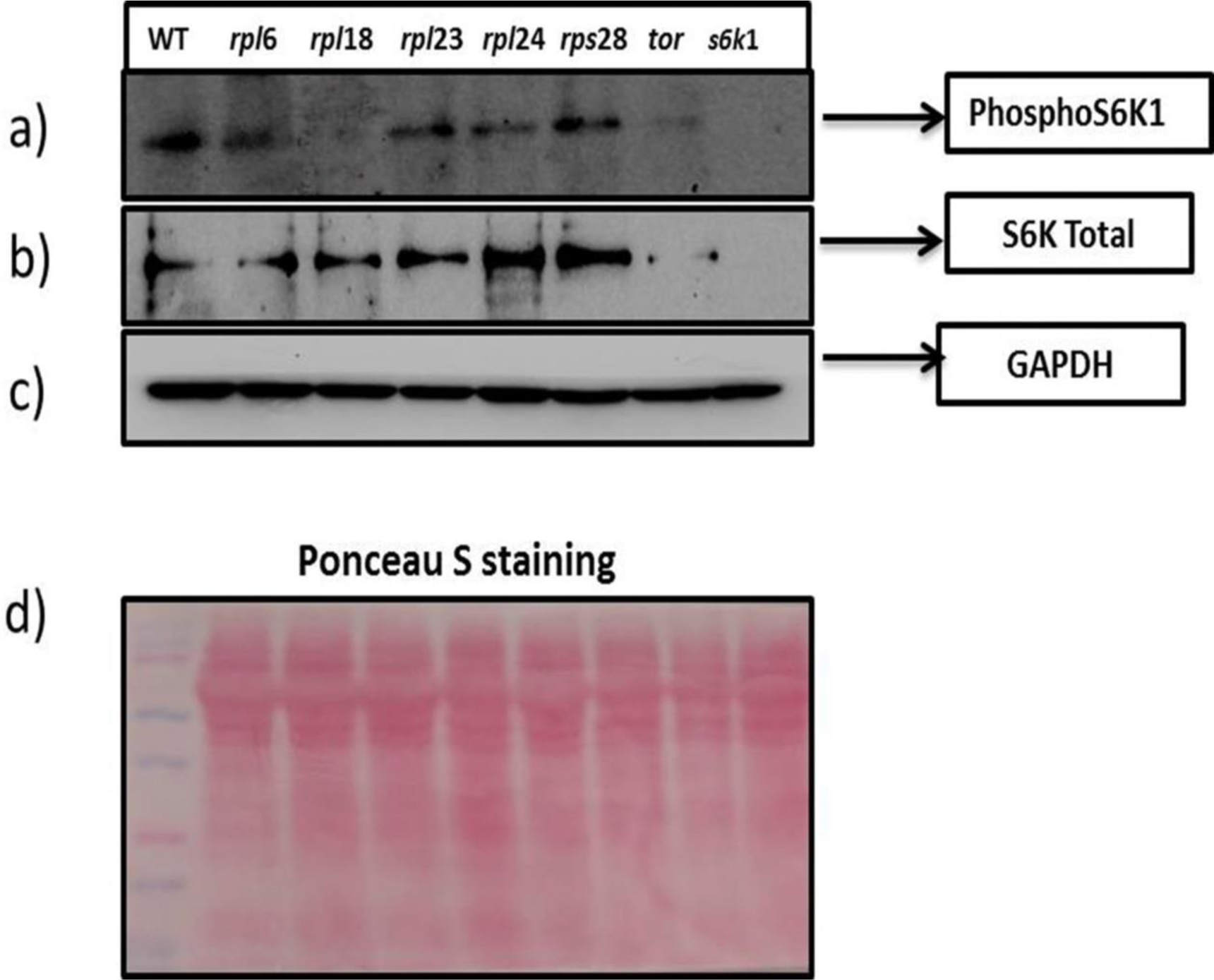
S6K1 phosphorylation assay in Arabidopsis T-DNA insertional mutants. Phosphorylation of p70kDa S6K1 at Thr389 residue was detected in Arabidopsis T-DNA insertional mutants of *tor*, *s6k*1 protein along with mutants of ribosomal proteins *rpl6*. *rpl18, rpl23a, rpl24, rps28a* and total protein isolated from WT Col0 Arabidopsis was taken as control. (a) phospho S6K1 detection in all mutants (b) western blot of total S6K protein (c) GAPDH protein used as loading control.

### 3.9. Identification of phosphorylation sites and protein kinase binding motifs in RPL and RPS proteins

The Ribosomal S6 protein kinases (RSKs) family proteins are Serine/ Threonine kinases that regulate cell growth, and proliferation. The two subfamilies of RSKs, p90 RSK and p70 S6K phosphorylate RPS6 at Ser/Thr residues for modulation of ribosome biogenesis and protein translation. The sequences of ribosomal proteins and their variants present in Arabidopsis and Rice genomes were analyzed for the presence of phosphorylation sites on Serine and Threonine residues by RSKs and their location on the protein (Table.1; Fig. 11a, b, c, d, e, f). The presence of phosphorylation sites suggests the possible interaction of the kinase with the RPs at the appropriate positions. The identified peptide sequences with the Ser/ Thr phosphorylation sites were then compared with the conserved RPS6 peptide sequences in Arabidopsis. The sequence alignment of *Oryza sativa* RPS6A/B proteins from ssp. *japonica* share highest similarity with *Arabidopsis thaliana* RPS6A/B proteins in their Ser/ Thr phosphorylation sites at similar amino acid positions (Thr 9, Thr 10, Thr 69, Thr 81, Thr 127, Thr 129, Thr 161, Thr 165, Thr 167, Thr 168, Thr 185, Thr 188, Thr 248, Thr 249, Ser33, Ser 37, Ser 98, Ser 105, Ser 109, Ser 141, Ser 150, Ser 175, Ser 208, Ser 229, Ser 231, Ser 237, Ser 240, Ser 241; Fig. 11a), The OsRPL6 protein has twenty five Ser/ Thr phosphorylation sites for various AGC kinases (PKA, PKB, PKC, PKG and RSK) when compared with the AtRPL6A/B proteins having twenty phosphorylation sites and exhibited maximum variation in their phosphorylation sites and their amino acid sequence adjacent to Ser/ Thr sites (Fig. 11b). A conserved Guanine residue is present in the peptide sequence of AtRPS6 at Thr 81 residue (RGTP) and at Thr 93 (PGTV) of AtRPL6 protein. Similarly, a conserved Leucine was observed at Ser 119 (QLSL) of AtRPL6 and Ser 109 (DLSV) of AtRPS6. The presence of conserved amino acids adjacent to the phosphorylation sites at Ser/ Thr residues in AtRPS6 and other RPs indicates their interaction with RSK (S6K1/2) proteins. The other ribosomal proteins consisted of similar conserved sites such as AtRPS6-Thr 91(RTGE)/ AtRPL18-Thr 72(MTGK), AtRPS6-Thr 129 (DTEK)/ AtRPL18-Thr 105(FTER). The results of sequence alignment of RPs with AtRPS6 also showed replacement of Ser or Thr to Thr or Ser phosphorylation sites with similar peptide sequences (AtRPS6-Ser105-VSPDL to AtRPL18-Thr 86-ITDDL). Similarly, the Thr 81 of AtRPS6 (LHRGT) is replaced by Ser in AtRPL24 and OsRPL24 with peptide sequences LFLNS and LFANS respectively. The above results suggested that the RSKs possibly phosphorylate and activate other RPs in the same manner as they phosphorylate RPS6 protein.

## 4. Discussion

Plant TOR signaling has been investigated for its cross-links with several cellular functions, but its function in stress tolerance is still far from clear. In this context, we attempted to investigate the roles of TOR signaling in response to various abiotic stresses in Arabidopsis using TOR overexpression lines in this study. A reduction in TOR kinase activity caused by environmental stresses results in changes in carbon and nitrogen metabolism (Caldana et al. 2013) suggesting the role of TOR signaling mediated through nutrient sensing switch for regulation of metabolic processes (Mayordomo et al. 2002; Matsuo et al. 2007) The advancement in the knowledge of plant TOR signaling would present new insights into the understanding of stress tolerance mechanisms in plants. The inhibition of TOR signaling in plants might also create changes in cellular metabolism and accumulation of amino acids (Laplante and Sabatini, 2012; Moreau et al. 2012). TOR signaling modulates protein synthesis and ribosome biogenesis for the supply of uninterrupted energy and nutrient availability, which are essential for sustaining plant growth under stressful environments (Wullschleger et al. 2006; Ren et al. 2011). We observed abiotic stress tolerance and high biomass in terms of enhanced shoot and root growth in both *TOR*-OE Arabidopsis lines used in the present study and the enhanced other phenotypes associated with increased yield traits and water-use efficiency in rice lines that expressed AtTOR ectopically (Bakshi et al. 2017) reported by us earlier.

Salt and osmotic stresses are common abiotic factors that exhibit an overlap in their signaling pathways and have a negative influence on plant growth and development. Also, these stress conditions modulate the transcriptional and translational status of the cell. The phosphorylation of S6K1, the downstream target of TOR is sensitive to osmotic stress and this sensitivity is relieved by the co-expression of RAPTOR1 (Mahfouz et al. 2006). The overexpression of S6K1 conferred hypersensitivity to 5% Mannitol in Arabidopsis plants (Mahfouz et al. 2006). The constitutive expression lines of *AtTOR* in rice also exhibited upregulation of TORC1 components such as *RAPTOR* and *LST8*. This suggests that their overexpression results in maintenance of balanced transcript levels of *TOR* (which is usually inactivated under stress conditions). This balanced TORC1 signaling in response to stress occurs through the activity of RAPTOR1 (Mahfouz et al. 2006). Also, the *TOR* overexpression lines of Arabidopsis treated with salt (0.16 M NaCl) and osmotic (0.35 M mannitol) stress for 6–8 h in liquid ½ MS medium had no effect on oxidative stress induced autophagy suggesting that the TOR activation stabilizes the detrimental effect of stress conditions in plants (Pu et al. 2017). In one of our previous studies, the *AtTOR* in rice transgenic plants evidenced increased transcript level of stress inducible genes (Bakshi et al. 2017). To understand the role of TOR in response to long term exposure of various abiotic stresses in native system, the 15 DAG plants of two *TOR*-OE Arabidopsis lines were subjected to osmotic and salt stress treatments.

The TOR signaling reprograms the energy supply in the cells, which is lost during stress conditions. When we expressed the *TOR* constitutively in Arabidopsis, we observed that the transcript levels of *SOS*1 were upregulated 10-fold in the high and medium *TOR*-OE lines, ATR-1.4.27 and ATR-3.7.32 respectively in their treatment with high sodium chloride concentration. This indicates that the TOR signaling plays an important role in salt homeostasis by modulating the expression of appropriate protein(s) to protect from the detrimental activities of high salt concentration in cells. Ascorbate peroxidases play a key role in maintaining cellular redox homeostasis by ROS scavenging under drought and salinity stress (Asada 1992; Shigeoka et al. 2002; Mittler et al. 2004). The increased transcript level of ascorbate peroxidase in the two *TOR*-OE Arabidopsis lines ATR-1.4.27 and ATR-3.7.32 in all the abiotic stress treatments was also an additional evidence to state that the TOR pathway is modulates ROS signaling. TOR inhibition causes reduction in ROS signaling in roots, whereas the ROS accumulation is certainly important for root hair development (Foreman et al. 2003; Livanos et al. 2012). The reduced ROS levels in TOR suppressed FKBP12 overexpressing Arabidopsis lines were observed with phenotypes having low photosynthetic efficiency and retarded leaf development (Ren et al. 2012). The increased transcription of ascorbate peroxidase (*APX*1) gene in *TOR*-OE transgenic lines might help detoxification of stress induced hydrogen peroxide levels and this might help sustain the plant under stressful environments. The *cis-*acting ERF5 protein binds to the GCC or DRE/CRT motifs in response to osmotic stress. *ERF*5 gene was induced in *TOR*-OE lines in all stress treatments, which indicates the involvement of TOR signaling in ethylene mediated regulation of abiotic stress responsive genes (Lee et al. 2004; Wang et al. 2004; Zhang et al. 2009; Hussain et al. 2011). In addition, the disruption of TOR resulted in increased accumulation of secondary metabolites, triacylglycerides, amino acids and the tri carboxylic acid intermediates (Caldana et al. 2013; Li et al. 2015). The amino acid-derived secondary metabolite precursors, polyamines are important in providing tolerance to heat, drought, salinity and cold stresses (Gill and Tuteja 2010). The expression of SAMDC gene involved in polyamine biosynthesis was induced in the *TOR*-OE Arabidopsis transgenic lines under all stress treatments, suggesting that the elevated levels of polyamine biosynthesis would be helpful in countering deleterious effects of the stress treatments.

TOR is a central regulator of ribosome biogenesis and translation in eukaryotes. TOR phosphorylates p70kDa ribosomal S6 kinase (S6K)-1 at Thr-389 residue in animals and plants (Mahfouz et al. 2006; Xiong and Sheen 2012; Schepetilnikov et al. 2013; Fonseca et al. 2014; Ahn et al. 2015; Xiong et al. 2017). TOR-S6K1-RPS6 phosphorylation cascade has been demonstrated in Arabidopsis, rice, potato and other plant species (Ren et al. 2011; Xiong et al. 2013; Song et al. 2017). The TOR pathway is also linked to the transcriptional and translational regulation of RPL and RPS in mammals and yeast (Jorgensen et al. 2004; Kaeberlein et al. 2005; Moreau et al. 2012; Ren et al. 2012; Xiong et al. 2013; Fonseca et al. 2015). In turn, the ribosomal proteins are also interlinked in regulation of S6K phosphorylation and translational initiation. The ribosomal proteins such as AtRPS6A and AtRPS6B are equally important as TOR for plant growth and development particularly for embryonic viability and the corresponding knock out mutants showed perturbed phenotype in Arabidopsis (Ren et al. 2012). Both TOR and RPS6A or B act in a dose-dependent manner and also the overexpression lines of *RPS6* revealed phenotypes similar to *TOR*-OE lines (Ren et al. 2011; Ren et al. 2012; Bakshi et al. 2017). The extra-ribosomal functions of RPs in biotic and abiotic stress response have been demonstrated in rice (Moin et al. 2016; Moin et al. 2017; Saha et al. 2017). Co-immunoprecipitation assay predicted RPS3, RPS6, RPS7, RPS10, RPS11, RPS17 and RPL13A, RPL18, RPL18A, RPL19 and RPL23 as S6K interacting proteins with conserved phosphorylation sites, RXRXXT/S (Pavan et al. 2016). Similarly, mutation or loss of RPS19 and other RPs induce S6K phosphorylation with an increase in ROS (Reactive Oxygen Species) levels in zebrafish (Heijnen et al. 2014) and RPS27L silencing also led to autophagy in mouse fibroblasts and human breast cancer cells by inhibition of S6K1 phosphorylation and mTOR Complex1 activity (Xiong et al. 2018). However, these studies were conducted on animals and analogous data are not available in plant systems. Although, S6K1 phosphorylation might also be affected by the several other protein kinases (PKA or PKB), its regulation is variable in plant system and the exact mechanism of this is still not known. The S6K1 phosphorylation is reduced in Arabidopsis in response to osmotic stress (Mahfouz et al. 2006). In contrast to this, cold stress strongly up-regulated *AtS6K1* and *AtS6K2* transcripts in Arabidopsis (Mizoguchi et al. 1995).

The TOR-S6K1-RP signaling and protein synthesis are complex processes consisting of many unknown substrates and regulators or effectors in plants system. In this work, we have also addressed the qualitative changes in phosphorylation of S6K1 protein in Arabidopsis plants with loss of RP function. A PI3K (Phosphoinositide 3-Kinase) inhibitor, LY294002, efficiently suppressed translation and phosphorylation of phytohormone-induced *RPS6* and *RPS18A* mRNAs in Arabidopsis without affecting global translation in cell (Turck et al. 2004). The inactivation of TOR inArabidopsis TOR-RNAi lines also showed induced RPL13 and RPS14 proteins expression (Dobrenel et al., 2016). The studies on S6K-RPs interaction have been well demonstrated in animal system. In Animal cells, RPS6 is associated with mRNAs of 5’-TOP tract such as *RPL11* and *RPS16* and negatively regulates their translation (Hagner et al. 2011). The association of ribosomal proteins such as RPL6, RPL18, RPL24 along with other RPs has been reported as an essential step in translational transactivation in plants and animals with viral infection (Leh et al. 2000; Wang et al. 2002; Martínez and Daròs 2014; Li et al. 2018). These reports in plants and animals suggest the importance of extra-ribosomal association of RPs with RNAs and other proteins and also provided a link for the possible interaction with the S6K1 for translational control. *Bacillus subtilis*, a bacterium reportedly requires an essential binding of RPL6 with a GTPase (RbgA) for assembly of ribosome large subunit (Gulati et al. 2014). We further suggest a similar requirement of interaction of ATPase and RPs in regulating S6K1 phosphorylation in plants. RPS6 interaction with rRNA gene promoter is also reported in Arabidopsis (Kim et al. 2014). There are no supporting reports for the modulatory effect of RPs on TOR-S6K1 signaling pathway in plants (Ren et al. 2011; Kim et al. 2014). In our study, the loss of ribosomal proteins in Arabidopsis also differentially affected phosphorylation of S6K1 at Thr-389 residue. However, more work will be required to understand the mechanisms and function of RPs in S6K1 phosphorylation and its effect on plant translation (Fig. 12). The results from present work suggest the possible crosslink of TOR signaling with ribosomal proteins in feedback regulation of S6K1 and the possible involvement of this TOR-S6K1-RP cyclic regulation in providing tolerance to abiotic stress in plants.

**Figure 11.**
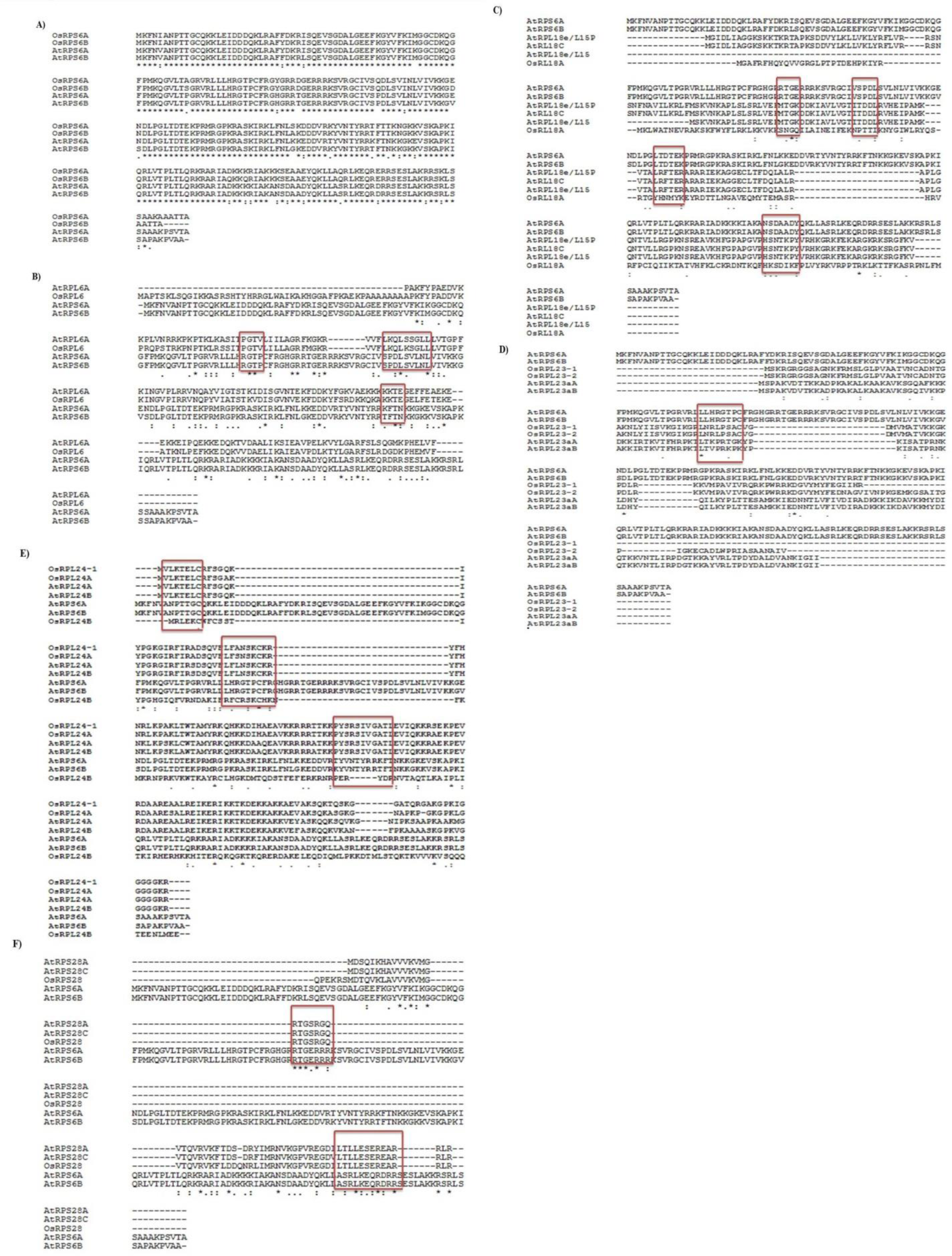
Sequence alignment of RPL and RPS protein sequences identifies conserved ser/ thr phosphorylation sites. The RPL and RPS protein sequences of Arabidopsis and rice were aligned using CLUSTAL Omega to identify the conserved sites for phosphorylation by AGC kinase family kinases specifically at Ser/ Thr residues. a) Sequence alignment of RPS6A/B proteins of Arabidopsis and oryza sativa ssp. indica, b) alignment of RPS6A/B proteins and RPL6, c) alignment of RPS6A/B proteins and RPL18, d) alignment of RPS6A/B proteins and RPL23, e) alignment of RPS6A/B proteins and RPL24, f) alignment of RPS6A/B proteins and RPS28,

**Figure 12.**
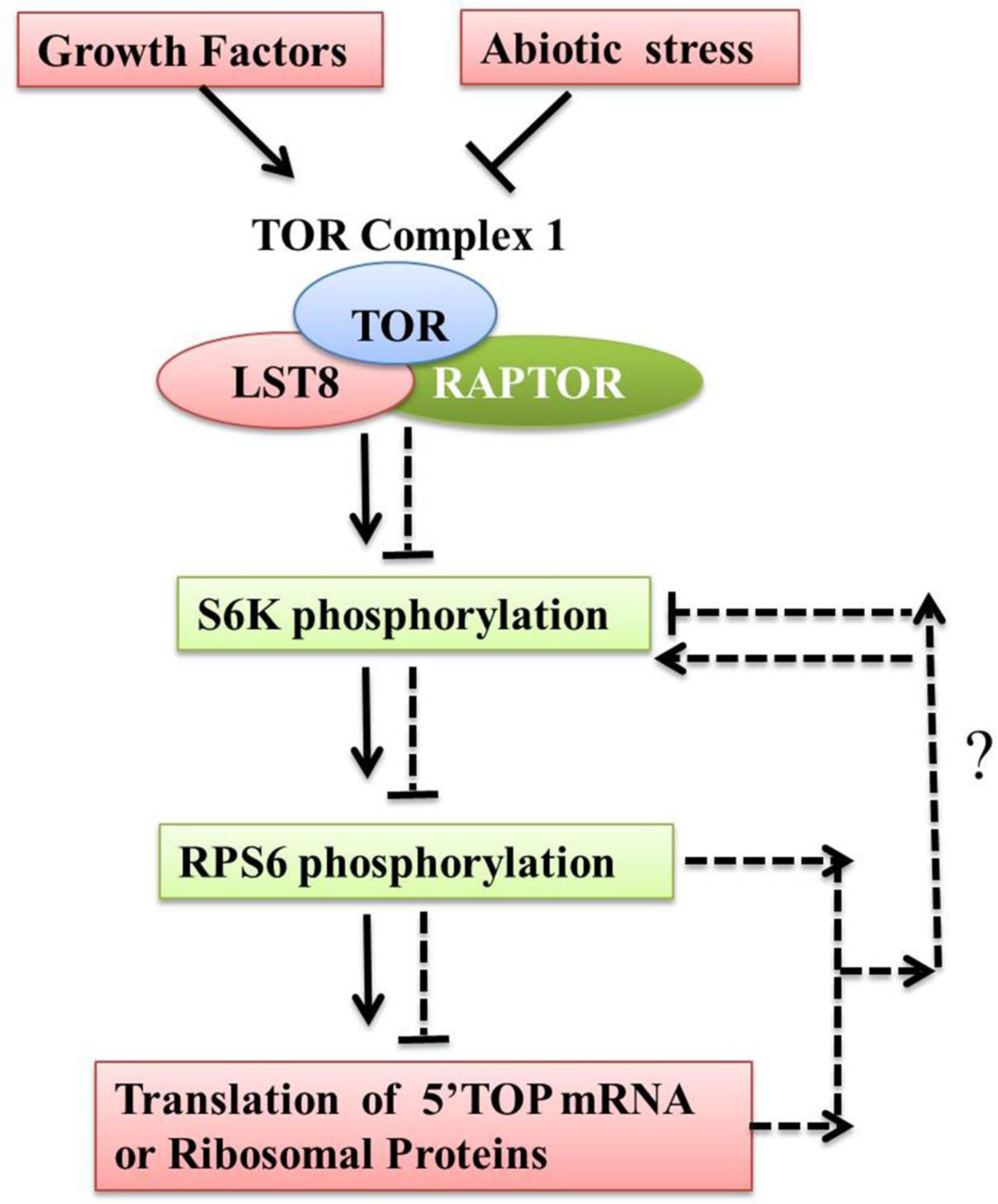
Possible feedback regulation of S6K1 phosphorylation by TOR pathway. TOR Complex 1 mediated phosphorylation of S6K1 regulates translational initiation by further phosphorylating RPS6 protein and the regulation of other RPL and RPS has been indirectly linked with the TOR pathways. In our observations, the inhibition of RPs and TOR differentially regulates S6K1 protein phosphorylation. Possibly the TOR and RPs are interlinked for regulation of S6K1 phosphorylation, where RPs also have an independent role in differentially regulating the S6K phosphorylation and modulate protein translation in plant cell. The figure represents a model for regulation of S6K1 phosphorylation by loss of RPs in plants, which is possibly mediated via association of RPs with the S6K1 protein or the 5’TOP mRNA or the other regulatory proteins in the TOR pathway. The dotted ‘T’ shaped bars and dotted arrows represent possible negative and positive regulation respectively.

### 4.1. Conclusion

TOR as a central coordinator regulates myriads of signaling cascades including the stress induced signaling. Previous findings showed the S6K1 phosphorylation is TORC1 dependent (Xiong & Sheen, 2012). This study indicated that the phosphorylation of S6K1 is also dependent on RPL and RPS protein function. The presence of Ser/ Thr phosphorylation sites in RPL and RPS proteins favors their possible interaction with the RSK protein kinases. Our study shows that the TOR pathway is also linked with the phosphorylation and activation of other RPs apart from the TOR-S6K-RPS6 signaling in plants. Western blot analysis in the present manuscript also is in line with the influence of other RPLs and RPS proteins whose mutation might reflect in the feedback regulation of S6K1 phosphorylation at Thr389 residue suggesting the possible interaction between the other RPs and S6K1 protein. In summary, the data presented in the study provides a resource for subsequent elucidation of S6K1-RPL/RPS proteins interaction and can be used in future experiments for the detection of the S6K1 protein activity in plants.

## AUTHOR CONTRIBUTION STATEMENT

AB, RD and PBK designed the work. AB performed all the experiments and analyses. RD provided the AtTOR vector. ABMR and MBG helped in S6K1 phosphorylation. AB, MM, MSM and PBK prepared the manuscript. All the authors read and approved the manuscript.

## ACKNOWLEDGEMENTS

AB acknowledges financial support from Department of Biotechnology-Research Associate program in Biotechnology and Life Sciences (2-29/RA/Bio/2018/550) and Department of Biotechnology, ICAR-Indian Institute of Rice Research (IIRR), Hyderabad. Authors acknowledge Dr. Gassmann Walter of Christopher S. Bond Life Sciences Center, University of Missouri for providing Arabidopsis SALK mutant lines of RPs and TOR. PBK acknowledges the National Academy of Sciences, India for the grant of Platinum Jubilee Senior Scientist.

