## Supplementary file for "Target of Rapamycin: function in abiotic stress tolerance in Arabidopsis and its involvement in a possible cross-talk with ribosomal proteins"

[5]Present address: Agri Biotech Foundation, Agricultural University Campus, Rajendranagar, Hyderabad-500030, Telangana, India

*** Correspondence to:**

**Contact details of authors:**

Mazahar Moin:;

M. S. Madhav:;

Meher B. Gayatri:;

Aramati B. M. Reddy:;

Raju Datla:

**Key words:** TOR, Abiotic stress, Ribosomal proteins Large and Small subunit genes

**Abbreviations:** *AtTOR*, *Arabidopsis thaliana Target of Rapamycin*; *TOR*-OE, *TOR* overexpressing; RPL, Ribosomal protein large subunit; RPS, Ribosomal protein small subunit; RPs, Ribosomal Proteins; TFs, Transcription factors; WT, Wild Type; DAG, Days after Germination; PEG, Polyethylene glycol; RSK, Ribosomal S6 kinase

**Fig 1.**

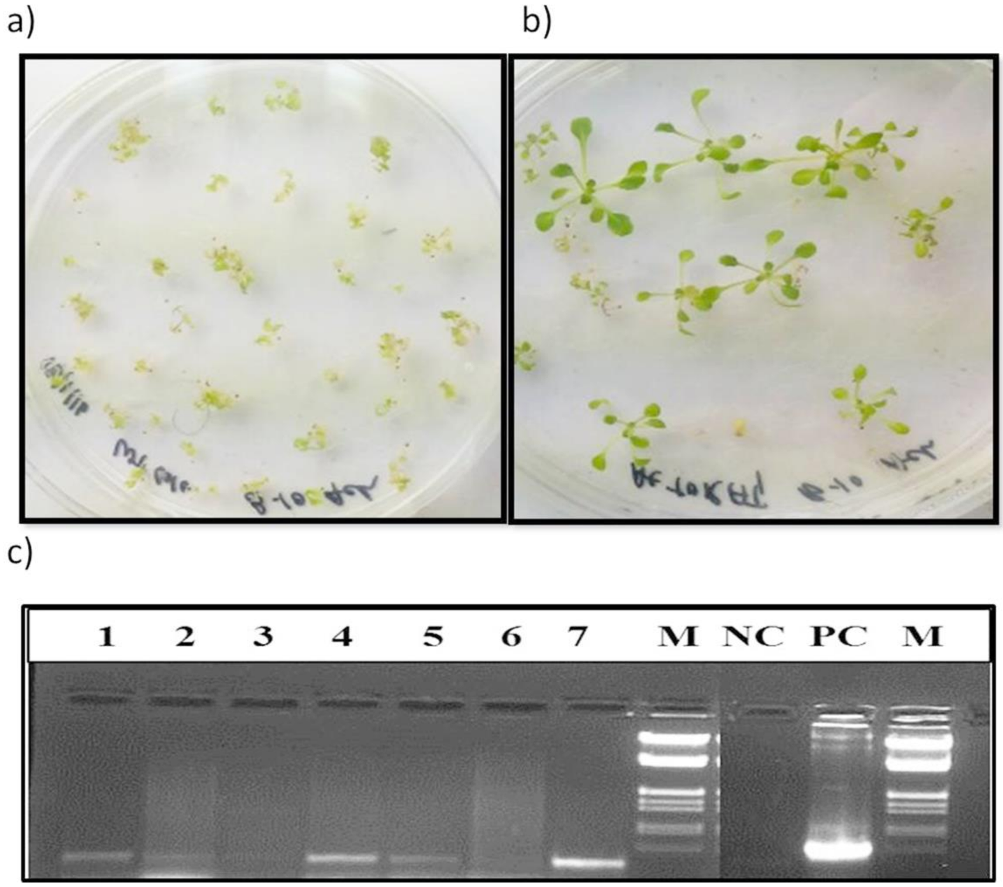


**Fig 1. Screening of transformed Arabidopsis plants in $T_1$ generation**

The seeds obtained from Agrobacterium infected primary plants were selected on MS medium containing Phosphinothricin (PPT, 10 mg $ml^{-1}$). (a) Germination of WT Arabidopsis (Col 0) seeds on PPT. (b) Germination of T1 generation seeds on PPT. (c) PCR screening of transformants selected on PPT medium with bar gene (550 bp) in $T_1$ generation. M, λ.*Eco*RI-*Hind*III DNA Marker; PC, Positive Control; NC, Negative Control, 1-7 transgenic plants.

**Fig 2.**

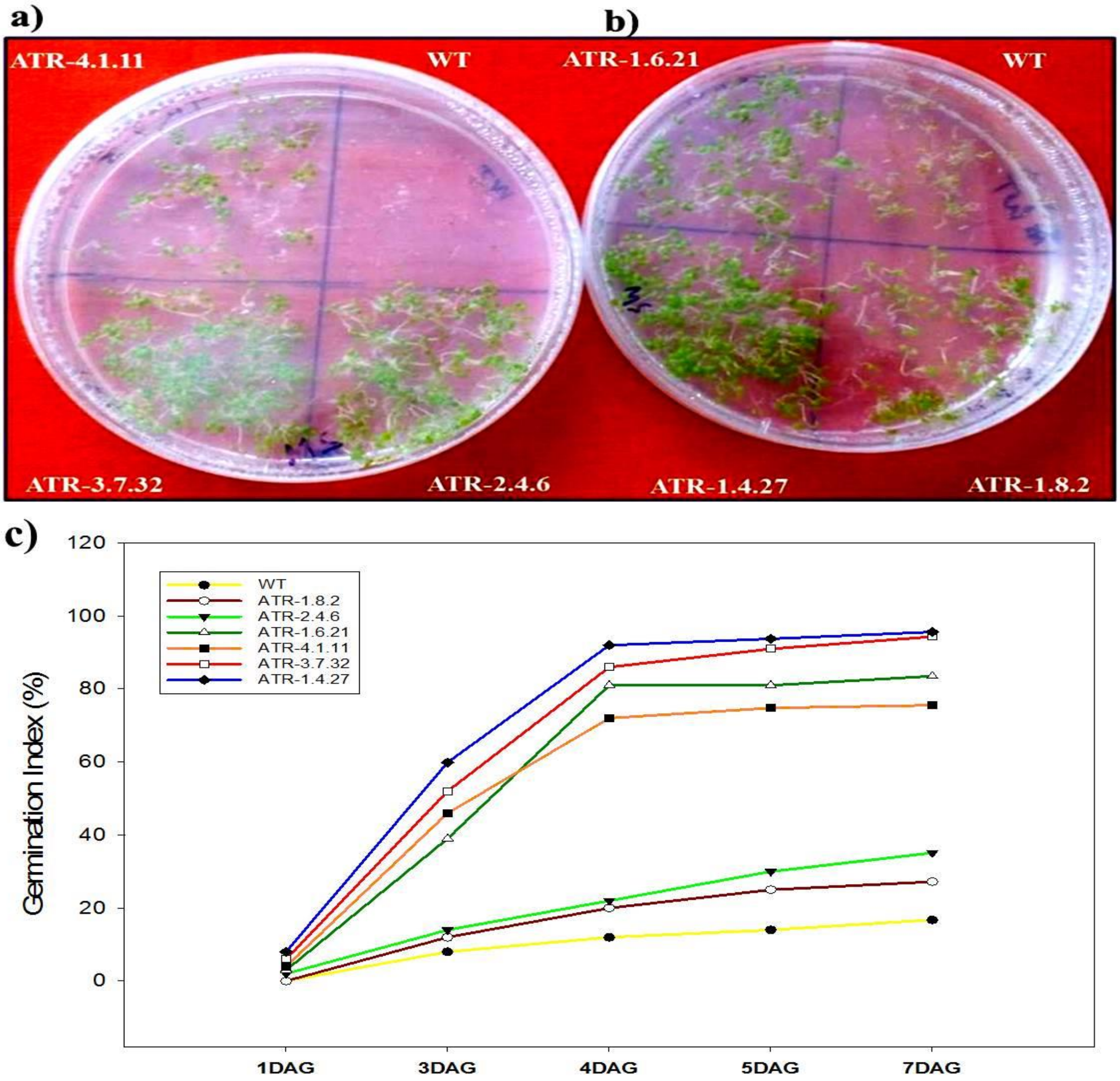


**Fig 2. Germination and screening of Arabidopsis $T_2$-transgenic seeds on selection medium**

The seeds obtained from PPT selected $T_1$ generation plants were further germinated on selection medium (containing ½ MS salts and 10 mg $ml^{-1}$, PPT) and the germination rate was calculated. (a, and b) The transgenic lines, ATR-1.4.27 and ATR-3.7.32 almost had 100% germination rate suggesting that they are homozygous in nature, whereas the lines, ATR-1.6.21 and ATR-4.1.11 had nearly 80% and the lines, ATR-1.8.2 and ATR-2.4.6 had ~20% germination rate suggesting their hemizygous or heterozygous nature. The WT Arabidopsis seeds became bleached on PPT selection medium. (c) The graphical representation of seed germination rate (%) in selected Arabidopsis transgenic lines in $T_2$ generation.

**Fig 3.**

a)

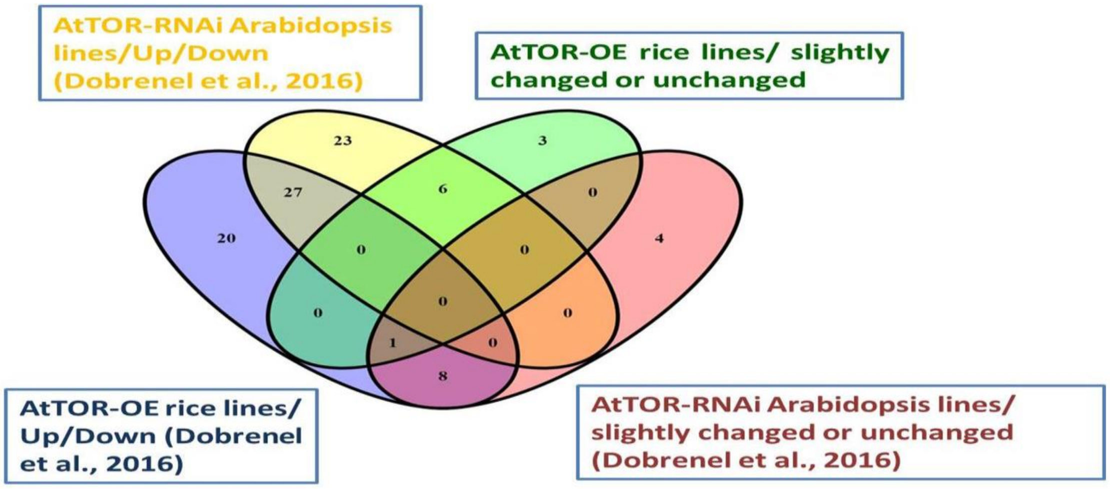


b)

| Change in Expression | TOR gene overexpression or RNAi inactivation | |
|---|---|---|
| | AtTOR-OE, TR-2.24/ TR-15.1 rice lines used in present study | AtRNAi Arabidopsis lines observed by (Dobrenel et al., 2016, given as the log2 value =+1 showed as upregulation) |
| TOR overexpression or inactivation resulted in change in the RP genes expression either Up/Down | RPL4, RPL6, RPL7, RPL8, RPL10, RPL11, RPL13a, RPL18p, RPL18a, RPL19.3, RPL21, RPL24b, RPL26, RPL28, RPL29, RPL30, RPL31, RPL32, RPL34, RPL35a, RPL36.2, RPL38a, RPL44, RPL51, RPS3a, RPS3-1, RPS4, RPS4a, RPS5, RPS5a, RPS6, RPS6a, RPS7, RPS7a, RPS9, RPS9-2, RPS10, RPS10a, RPS13, RPS13a, RPS15, RPS15a, RPS17, RPS19, RPS20, RPS21, RPS23, RPS24a, RPS25, RPS25a, RPS26, RPS27, RPS27a, RPS28, RPS29, RPS30 | RPL3, RPL4, RPL5, RPL6, RPL7a, RPL8, RPL9, RPL10. RPL10a, RPL11, RPL12, RPL14, RPL15, RPL18, RPL18a, RPL19, RPL21, RPL22, RPL23, RPL23a, RPL24, RPL27, RPL27a, RPL28, RPL30, RPL31, RPL32, RPL34, RPL35, RPL35a, RPL36, RPL36a, RPL37, RPL37a, RPL38, RPL39, RPL40, RPL41, RPS2, RPS3, RPS6, RPS9, RPS10, RPS11, RPS12, RPS13, RPS15, RPS16, RPS17, RPS18, RPS19, RPS21, RPS26, RPS27, RPS27a, RPS28 |
| TOR overexpression or inactivation resulted in change in the RP genes expression Slightly Up/Unchanged | RPL5, RPL15, RPL23, RPL24, RPL27, RPL37, RPS18a, RPS18b, RPS23, RPS24 | RPL7, RPL13, RPL13a, RPL17, RPL26, RPL29, RPS5, RPS8, RPS14, RPS20, RPS23, RPS29, RPS30 |

**Fig 3. Comparison of expression of RPL and RPS genes in *AtTOR* overexpressing rice lines and *AtTOR*-RNAi Arabidopsis lines**

**a, b)** The expression analysis data of RPL and RPS genes in the two high *AtTOR* expressing rice transgenic lines TR-2.24 and TR-15.1 (Bakshi et al., 2017) was compared with the transcriptomic data of cytoplasmic RPL and RPS genes in two AtTOR-RNAi lines obtained from Dobrenel et al., (2016). More than 35 genes were highly upregulated or changed in their expression in both rice and Arabidopsis lines, whereas more than ten genes were slightly changed or remain unchanged in the two systems. These results suggested that TOR acts differently in regulation of RP genes transcription when overexpressed or inhibited. The TOR inactivation and RP genes upregulation also supported that the increased RP transcript level is linked to the TOR inhibition under abiotic stress conditions.

**Table 1. List of primers used in analysis of transgenic plants**

| S. No. | Primer Name | Sequence (5'-3') |
|---|---|---|
| 1. | Bar FP<br>Bar RP | GAATTCATGAGCCCAGAACGACGC<br>CCCGTCACCGAGATCTGAAAGCTT |
| 2. | AtTOR FP<br>AtTOR RP | GCGGCCGCATGTCTACCTCGTCGCAATC<br>CCCGGGTGAGGATCCAAAGCGCCCATAAT |
| 3. | CaMV35S FP<br>CaMV35S RP | AAGCTTGATCCCCAACATGGTGGAGC<br>TTCGGATCTAGATATCACATCAATC |
| 4. | OsTOR RT FP<br>OsTOR RT RP | CAAATCGTATGGGAGGAGCTA<br>GCAGCCATAAGAAGTTTCTCCA |
| 5. | AtTOR RT FP<br>AtTOR RT RP | ATTTCTTCTGCGATCTTCGG<br>CAACACTTGGGCAGAGAGAG |
| 6. | OsActin1RT FP<br>OsActin1RT RP | TCCCCCATGCTATCCTTCG<br>TGAATGAGTAACCACGCTC |
| 7. | OsTubulinRTFP<br>OsTubulinRTRP | TGACCACACCTAGCTTTGG<br>AGGGAACCTTAGGCAGCATG |
| 8. | AtActin2RT FP<br>AtActin2RT RP | CAGCAGATGTGGATCTCCAAGG<br>CGCAGACGTAAGTAAAAACCC |

**Table 2: Transmission of T-DNA to $T_2$ plants (∑ Chi Square value ($\chi 2$) Segregation Analysis).**

| S. No. | Trangenic Line Name | Total number of seeds screened | Number of PPT positive (Germinated seeds) | Number of PPT negative (Non-Germinated seeds) | ∑ Chi Square value ($\chi 2$) | Germination Percentage on PPT (%) | Transgene Expression Level |
|---|---|---|---|---|---|---|---|

| | | | | | | | |
|---|---|---|---|---|---|---|---|
| **1** | ATR-1.2.23 | 97 | 78 | 19 | 0.378865979 | 80.4 | high |
| **2** | ATR-1.4.12 | 98 | 76 | 22 | 0.085034014 | 77.6 | high |
| **3** | ATR-1.4.27 | 104 | 99 | 5 | 5.653846154 | 95.2 | high |
| **4** | ATR-1.6.21 | 85 | 48 | 37 | 3.891176471 | 56.5 | high |
| **5** | ATR-1.7.10 | 95 | 78 | 17 | 0.639473684 | 82.1 | high |
| **6** | ATR-1.8.2 | 93 | 44 | 49 | 9.506272401 | 47.3 | low |
| **7** | ATR-2.4.6 | 91 | 61 | 30 | 0.77014652 | 67.1 | medium |
| **8** | ATR-2.4.28 | 109 | 77 | 32 | 0.275993884 | 70.6 | medium |
| **9** | ATR-3.5.42 | 114 | 88 | 26 | 0.073099415 | 77.2 | high |
| **10** | ATR-3.6.24 | 105 | 81 | 24 | 0.064285714 | 77.1 | high |
| **11** | ATR-3.7.32 | 106 | 97 | 9 | 3.852201258 | 91.5 | medium |
| **12** | ATR-3.8.41 | 104 | 89 | 15 | 1.551282051 | 85.6 | medium |
| **13** | ATR-4.1.11 | 93 | 42 | 51 | 11.04032258 | 45.2 | low |

**Table 3. List of Primers used in qRT-PCR analysis of RPS genes**

| **S. No.** | **Primer name** | **Sequence 5'- 3'** |
|---|---|---|
| 1. | S3a RT FP<br>S3a RT RP | TGAAGCCCATGTGGATGTTA<br>TCACATGTTGATGCCTGGTT |
| 2. | S4 RT FP<br>S4 RT RP | CAGGTTGAAGTATGCGCTGA<br>AGCGACCCTTGGTGTCATAG |
| 3. | S4a RT FP<br>S4a RT RP | AGCGGCATGTTATGGTTGAT<br>GGACCTTGCAGAGCTTGAAC |
| 4. | S5 RT FP<br>S5 RT RP | ATATCCCTCGCCGACTACCT<br>TGATCTTCTTGCCGTTGTTG |
| 5. | S5a RT FP<br>S5a RT RP | CAACGGGAAGAAGATCATGG<br>GACCCTCCTCAAGGGAGAGA |
| 6. | S6 RT FP<br>S6 RT RP | GCTACGGCATCAAGAAGCTC<br>CAAGTGCCGTGTCACAAAGT |
| 7. | S6a RT FP<br>S6a RT RP | CATGAAGCAGGGTGTGCTTA<br>TCAGTCAAGCCAGGAAGGTC |

| | | |
|---|---|---|
| 8. | S7 RT FP<br>S7 RT RP | TGTGCAAGGCATTCAAGAAA<br>TCCAAGATTCCATCATGAACAG |
| 9. | S7a RT FP<br>S7a RT RP | AGAAGAAGTTCAGCGGCAAG<br>GCACCATCCAGACGGTATCT |
| 10. | S9 RT FP<br>S9 RT RP | TTATCAGGCAGCACCACATC<br>GGCCTTCTGGTTCTTCCTCT |
| 11. | S9-2 RT FP<br>S9-2 RT RP | GAGCTGTGGCGTGTTCAGTA<br>AACAGTGAGGGCAAGGACAT |
| 12. | S10 RT FP<br>S10 RT RP | CCCTCAAGAAGTCTGCCAAG<br>ACCAAAATCACCTGGAGCAC |
| 13. | S10a RT FP<br>S10a RT RP | GCATTGAGCACCTGAGGAAT<br>AACCTTGGTCTGTCCCCTTC |
| 14. | S13 RT FP<br>S13 RT RP | AGGTGGAGGAGATGATCGTG<br>GACGGCCTTCTTGATGAGG |
| 15. | S13a RT FP<br>S13a RT RP | GTGGTGCTCCGTGACCAG<br>TGGAGTCCTTGTCCTTCCTG |
| 16. | S15 RT FP<br>S15 RT RP | ATGACCTCGTCCAGCTCTTC<br>CCGATCATCTCAGGGACAAT |
| 17. | S15a RT FP<br>S15a RT RP | CCGACCCTCATCAAAGGTTA<br>TCCTTGACACCAACATCGAA |
| 18. | S17 RT FP<br>S17 RT RP | AACAAGAAGGTGCTGGAGGA<br>TCCTGGAGCTTGAGGGAGAT |
| 19. | S18 RT FP<br>S18 RT RP | CGACATCGACATGAACAAGC<br>TCAGCCTCTCAAGGTCATCC |
| 20. | S18a RT FP<br>S18a RT RP | CTCACCTCCATCAAGGGTGT<br>ACCTCCCGTCCTTGTAGTCC |
| 21. | S18b RT FP<br>S18b RT RP | AAGGATGGGAGGTTCTCTCAG<br>TCTTGGAGACACCGACAGTCT |
| 22. | S19 RT FP<br>S19 RT RP | CAAGATGGAGCTCCCTGAGT<br>GGACGTGAGCCATTCCTCT |
| 23. | S20 RT FP<br>S20 RT RP | GCACCAGGATGAAGAAGGTT<br>TGCGATGGTTACCAGTATTCC |

| | | |
|---|---|---|
| 24. | S21 RT FP<br>S21 RT RP | TCAGATGGTGGACCTCTACG<br>AGAGCACTGTCGGCGTCT |
| 25. | S23 RT FP<br>S23 RT RP | GAGCCATCTTGGCAATGAAT<br>CGATGAAGTTCAAGCAACCA |
| 26. | S23a RT FP<br>S23a RT RP | TTTTGGGAGAAAAAGGCTTG<br>CCTCTGCTTCTTCTTCTCCTTG |
| 27. | S24 RT FP<br>S24 RT RP | GTCTCCAAGGCTGAGCTGAA<br>CTTGGGCTCGTACTTCTTCG |
| 28. | S25 RT FP<br>S25 RT RP | GGGAAAGCAAAAGGAGAAGG<br>GATTAGGCCCCGTGTCATTA |
| 29. | S25a RT FP<br>S25a RT RP | CGGAGGCAAGCAGAAGAAG<br>CAAGATCCTTGATGGCCTGT |
| 30. | S26 RT FP<br>S26 RT RP | CAAGGCGATCAAGAGGTTTC<br>GACGATGTGAGCATGGATTG |
| 31. | S27 RT FP<br>S27 RT RP | CGGAGCTGGAGAAGCTCA<br>CTGGCCTTCCCACCAGTAG |
| 32. | S27a RT FP<br>S27a RT RP | CGAAGATCCAGGACAAGGAG<br>CTTGGGCTTGGTGTACGTCT |
| 33. | S28 RT FP<br>S28 RT RP | GGATACCCAGGTCAAACTTGC<br>CTGGCCTCCCTCTCAGACT |
| 34. | S29 RT FP<br>S29 RT RP | CACTCCAACGTGTGGAACTC<br>CGGTACTTGATGAAGCCAATG |
| 35. | S30 RT FP<br>S30 RT RP | GAAGGTGAGGGGGCAGAC<br>GACGAAACGGCGGTTGTACT |

**Table. 4. List of primers used in qRT-PCR analysis of RPL genes**

| **S. No.** | **Primer Name** | **Sequence (5'-3')** |
|---|---|---|
| 1. | RPL3 RT FP<br>RPL3 RT RP | TGGACTTGTGGCCTATGTGA<br>CCGGCATCGCTATCATACTT |
| 2. | RPL4 RT FP<br>RPL4 RT RP | AAGAAGCTCGACGAGGTGTA<br>CCACATTCTTCAGAGGGTTC |
| 3. | RPL5 RT FP<br>RPL5 RT RP | GATCTTGGCATCAAGTACGAC<br>GACACCCTCATACTTGACCTG |
| | RPL6 RT FP | GTTCCTCAAGCAGCTCAAAT |

| | | |
|---|---|---|
| 4. | RPL6 RT RP | CTTCTGCTTCTTGTCCCTAGA |
| 5. | RPL7 RT FP<br>RPL7 RT RP | TACCCAAACCTGAAGAGTGTC<br>GACAGTCATGATCTCGTGGA |
| 6. | RPL8 RT FP<br>RPL8 RT RP | ACTACGCCATCGTCATCAG<br>GGTACTTGTGGTAGGCGTTT |
| 7. | RPL10 RT FP<br>RPL10 RT RP | AGAAGAAGCCTGGATTAGAGC<br>ATATCCTGCTGGAGGACTTG |
| 8. | RPL11 RT FP<br>RPL11 RT RP | AAGAAGATCGGTGAGGACATC<br>TCTTGACCTTCTTCCTGTCC |
| 9. | RPL12 RT FP<br>RPL12 RT RP | GCTCATTTGTACAGCACAGAG<br>TTGGTTCAGTCTGAGAAGGAG |
| 10. | RPL13a RT FP<br>RPL13a RT RP | GAACTACCACGACACCATCAG<br>GGGGCCAAAATATCTATCTG |
| 11. | RPL13b RT FP<br>RPL13b RT RP | AAGCACTGGCAGAACTATGTC<br>CCCTCGACTTCATGTTGTACT |
| 12. | RPL14 RT FP<br>RPL14 RT RP | GTGAACTACGGCAAGGACTAC<br>TAACATCAGCCTCCTCCATAG |
| 13. | RPL15 RT FP<br>RPL15 RT RP | ACAAGTACGTGTCGGAGCTAT<br>GACACGGTAAACCACATAACC |
| 14. | RPL18a RT FP<br>RPL18a RT RP | TCCAAGTTCTGGTACTTCCTG<br>GTTGTGGTAACCTGTTCTGCT |
| 15. | RPL18p RT FP<br>RPL18p RT RP | TGGGGAGGACTACTATGTTGA<br>AAACCTCTTGTCACTGTGAGG |
| 16. | RPL19.3 RT FP<br>RPL19.3 RT RP | AGTATCGTGAGGCCAAGAAG<br>CTTAGCCTCAAACTGGTCAGA |
| 17. | RPL21.2 RT FP<br>RPL21.2 RT RP | CTGAGGAAGATCAAGAACGAC<br>AACCACCCTTGAGATCATTG |
| 18. | RPL22 RT FP<br>RPL22 RT RP | GAGGTGAAAGGTCTGGATGTT<br>TCACTGGTTCTTCCTTCTCTG |
| 19. | RPL23A RT FP<br>RPL23A RT RP | GACCAAAGACCCTGAAGAAGG<br>ACGATGAAGACAAGGGTGTTG |
| 20. | RPL24b RT FP<br>RPL24b RT RP | GTTGGTGCTACACTGGAAGTT<br>CCTTCGACTGTGTCTTCTGAG |

| | | |
|---|---|---|
| 21. | RPL26.1 RT FP<br>RPL26.1 RT RP | ACAAGTACAACGTGGTGAGG<br>GTCCTTGTCGAGCTTGAGTT |
| 22. | RPL27.3 RT FP<br>RPL27.3 RT RP | CTTCCTCAAGCTCGTCAACT<br>CTTGGTGAAGAACCACCTGT |
| 23. | RPL28 RT FP<br>RPL28 RT RP | TAGACGAATACCTCCTGAAGA<br>AAACCCTGTTCGATCTTAGTC |
| 24. | RPL29 RT FP<br>RPL29 RT RP | CCCAACAAGCTCTCCAATATA<br>AGAAACAGAAGCATTCCCTG |
| 25. | RPL30e RT FP<br>RPL30e RT RP | GAGCAAGAAGAAGAACAAGTC<br>GCTTCATCCATATCTTTTCCG |
| 26. | RPL31 RT FP<br>RPL31 RT RP | TCAAGGAGATCAGGAAGTTTG<br>AACAGTGACCAGAGAGTAGAG |
| 27. | RPL32 RT FP<br>RPL32 RT RP | GCCTAATATTGGCTATGGTTC<br>CTTCTTCGTTGAGACATTGTG |
| 28. | RPL34 RT FP<br>RPL34 RT RP | GAAGAAGATCCAGGGAATTCC<br>CACAATCTTCTGCTCTTCAAC |
| 29. | RPL35a.3 RT FP<br>RPL35a.3 RT RP | CTACGTCTACAAGGCCAAG<br>TGCTGGGGTACATGAAGA |
| 30. | RPL36.2 RT FP<br>RPL36.2 RT RP | GGAAAAGTACCAAGAGAGTGA<br>CTTCTTCTTTGCTCTCTTGTG |
| 31. | RPL37 RT FP<br>RPL37 RT RP | CTTCCACCTGCAGAAGAG<br>CCCCTCTCTGAAGTTACTCT |
| 32. | RPL38 RT FP<br>RPL38 RT RP | CACGAGATCAAGGACTTCC<br>AAAGGTGGATGAAATGTAGGC |
| 33. | RPL44 RT FP<br>RPL44 RT RP | AAGAAGACCTACTGCAAGAAC<br>CCTTACCCTTCTTGTACTGAG |
| 34. | RPL51 RT FP<br>RPL51 RT RP | GTGACAGAGTTAGTCCGTGGA<br>TCTCAGCTTCACCACTTTCCT |

**Table. 5. List of primers used in qRT-PCR analysis of stress-inducible genes in TOR-OE Arabidopsis lines.**

| No. | Primer Name | Sequence (5'-3') |
|---|---|---|
| | AtSOS1 RT FP | ATCACTCGCTGCATCCAACT |

| | | |
|---|---|---|
| 1. | AtSOS1 RT RP | GAACCAATGCACTTTCCTGCC |
| 2. | AtERF5 RT FP<br>AtERF5 RT RP | GACAAAGACCGTGGGGGAAA<br>TCTTCTCACCTTCATCGGCG |
| 3. | AtERD11 RT FP<br>AtERD11 RT RP | ACACACTCCCATTTGTGTATTTCTT<br>CCAAAGGGGTTGCGAAGGAT |
| 4. | AtAPX1 RT FP<br>AtAPX1 RT RP | TTCTGAGCTTGGGTTTGCTGA<br>GCAAAAGCGCAACGGATGT |
| 5. | AtSAMDC RT FP<br>AtSAMDC RT RP | CAGTCGTTTCCTCACCGTCA<br>ATAGGTCCTGCAGAGGCAGA |
| 6. | AtMnSOD RT FP<br>AtMnSOD RT RP | TCCCCTACGACTATGGAGCA<br>CGGTATGCAATTTGGCGACG |
| 7. | AtActin2 RT FP<br>AtActin2 RT RP | CCAGCAGATGTGGATCTCCAAGG<br>CGCAGACGTAAGTAAAAACCCAG |
| 8. | AtTubulin-α RT FP<br>AtTubulin-α RT RP | TGTGCTCATCTTGCCACCACGGA<br>CTCAAGAGGTTCTCAGCAGTA |

**Table. 6. Holms-Bonferroni adjustment of p-values of expression data after the stress treatment analysis to *TOR*-OE plants.**

| Treatment | Gene name | Sample name | p- value | alpha altered | alpha critical | Holms-Bonferroni correction |
|---|---|---|---|---|---|---|
| Untreated | AtERD11 | UntWT | 0.056 | 0.000468333 | 0.054662398 | Non-Significant |
| | | Unt7.32 | 0.0159 | 0.001325 | 0.0147093568 | Significant |
| | | Unt4.27 | 0.0194 | 0.001616667 | 0.0176471388 | Significant |
| NaCl | | NaCl Wt | 0.0333 | 0.002775 | 0.0283561518 | Significant |
| | | NaCl7.32 | 0.0394 | 0.003283333 | 0.0326082943 | Significant |
| | | NaCl4.27 | 0.0882 | 0.00735 | 0.0587392013 | Non-Significant |
| PEG | | PEGWT | 0.0333 | 0.002775 | 0.0283561518 | Significant |
| | | PEG7.32 | 0.0609 | 0.005075 | 0.045694821 | Significant |
| | | PEG4.27 | 0.0674 | 0.005616667 | 0.0491301581 | Significant |

| | | | | | | |
|---|---|---|---|---|---|---|
| **Sorbitol** | | **SorbiWT** | 0.0217 | 0.001808333 | 0.0195228136 | **Significant** |
| | | **Sorbi7.32** | 0.0316 | 0.002633333 | 0.0271244357 | **Significant** |
| | | **Sorbi4.27** | 0.054 | 0.0045 | 0.0417961486 | **Significant** |
| **Mannitol** | | **mannWT** | 0.0215 | 0.001791667 | 0.0193614074 | **Significant** |
| | | **mann7.32** | 0.0219 | 0.001825 | 0.0196838995 | **Significant** |
| | | **mann4.27** | 0.067 | 0.000558333 | 0.064822297 | **Non-Significant** |
| **Untreated** | **AtMSD1** | **UntWT** | 0.0666 | 0.000555 | 0.064447943 | **Non-Significant** |
| | | **Unt7.32** | 0.0025 | 2.08333E-05 | 0.002496904 | **Significant** |
| | | **Unt4.27** | 0.0035 | 2.91667E-05 | 0.003493933 | **Significant** |
| **NaCl** | | **NaCl Wt** | 0.0035 | 2.91667E-05 | 0.003493933 | **Significant** |
| | | **NaCl7.32** | 0.0055 | 4.58333E-05 | 0.005485028 | **Significant** |
| | | **NaCl4.27** | 0.065 | 0.000541667 | 0.062949039 | **Non-Significant** |
| **PEG** | | **PEGWT** | 0.055 | 0.000458333 | 0.053526785 | **Non-Significant** |
| | | **PEG7.32** | 0.0015 | 0.0000125 | 0.001498885 | **Significant** |
| | | **PEG4.27** | 0.0045 | 0.0000375 | 0.004489974 | **Significant** |
| **Sorbitol** | | **SorbiWT** | 0.0035 | 2.91667E-05 | 0.003493933 | **Significant** |
| | | **Sorbi7.32** | 0.0265 | 0.000220833 | 0.026154806 | **Significant** |
| | | **Sorbi4.27** | 0.0301 | 0.000250833 | 0.02965517 | **Significant** |
| **Mannitol** | | **mannWT** | 0.0054 | 0.000045 | 0.005385567 | **Significant** |
| | | **mann7.32** | 0.0406 | 0.000338333 | 0.039793458 | **Significant** |
| | | **mann4.27** | 0.065 | 0.000541667 | 0.062949039 | **Non-Significant** |
| **Untreated** | **AtAPX1** | **UntWT** | 0.0035 | 2.91667E-05 | 0.003493933 | **Significant** |
| | | **Unt7.32** | 0.006 | 0.0005 | 0.0058249597 | **Significant** |
| | | **Unt4.27** | 0.055 | 0.000458333 | 0.053526785 | **Significant** |
| **NaCl** | | **NaCl Wt** | 0.0045 | 0.0000375 | 0.004489974 | **Significant** |
| | | **NaCl7.32** | 0.065 | 0.000541667 | 0.062949039 | **Non-Significant** |
| | | **NaCl4.27** | 0.06 | 0.0005 | 0.058249597 | **Non-Significant** |
| **PEG** | | **PEGWT** | 0.007 | 0.000583333 | 0.0067625224 | **Significant** |
| | | **PEG7.32** | 0.005 | 4.16667E-05 | 0.004987624 | **Significant** |
| | | **PEG4.27** | 0.0045 | 0.0000375 | 0.004489974 | **Significant** |
| **Sorbitol** | | **SorbiWT** | 0.004 | 3.33333E-05 | 0.003992077 | **Significant** |
| | | **Sorbi7.32** | 0.06 | 0.0005 | 0.058249597 | **Non-Significant** |
| | | **Sorbi4.27** | 0.0041 | 3.41667E-05 | 0.004091676 | **Significant** |
| **Mannitol** | | **mannWT** | 0.005 | 4.16667E-05 | 0.004987624 | **Significant** |
| | | **mann7.32** | 0.0065 | 0.000541667 | 0.0062949039 | **Significant** |
| | | **mann4.27** | 0.07 | 0.000583333 | 0.067625224 | **Non-Significant** |
| **Untreated** | **AtSOS1** | **UntWT** | 0.0275 | 0.000229167 | 0.027128383 | **Significant** |
| | | **Unt7.32** | 0.0073 | 6.08333E-05 | 0.00727364 | **Significant** |

| | | | | | | |
|---|---|---|---|---|---|---|
| | | **Unt4.27** | 0.00757 | 0.000630833 | 0.0072927852 | **Significant** |
| **NaCl** | | **NaCl Wt** | 0.0124 | 0.000103333 | 0.01232407 | **Significant** |
| | | **NaCl7.32** | 0.0135 | 0.0001125 | 0.013410033 | **Significant** |
| | | **NaCl4.27** | 0.0207 | 0.0001725 | 0.020488975 | **Significant** |
| **PEG** | | **PEGWT** | 0.0217 | 0.000180833 | 0.021468169 | **Significant** |
| | | **PEG7.32** | 0.0265 | 0.000220833 | 0.026154806 | **Significant** |
| | | **PEG4.27** | 0.0299 | 0.000249167 | 0.029461033 | **Significant** |
| **Sorbitol** | | **SorbiWT** | 0.0302 | 0.000251667 | 0.029752224 | **Significant** |
| | | **Sorbi7.32** | 0.0361 | 0.000300833 | 0.035461404 | **Significant** |
| | | **Sorbi4.27** | 0.036 | 0.0003 | 0.035364917 | **Significant** |
| **Mannitol** | | **mannWT** | 0.0149 | 0.000124167 | 0.014790456 | **Significant** |
| | | **mann7.32** | 0.0164 | 0.000136667 | 0.016267355 | **Significant** |
| | | **mann4.27** | 0.0211 | 0.000175833 | 0.020880769 | **Significant** |
| **Untreated** | **AtSAMDC** | **UntWT** | 0.0376 | 0.000313333 | 0.036907571 | **Significant** |
| | | **Unt7.32** | 0.0043 | 3.58333E-05 | 0.004290845 | **Significant** |
| | | **Unt4.27** | 0.0418 | 0.000348333 | 0.04094541 | **Significant** |
| **NaCl** | | **NaCl Wt** | 0.0244 | 0.000203333 | 0.024107148 | **Significant** |
| | | **NaCl7.32** | 0.0479 | 0.000399167 | 0.04678001 | **Significant** |
| | | **NaCl4.27** | 0.0489 | 0.0004075 | 0.047733138 | **Significant** |
| **PEG** | | **PEGWT** | 0.049 | 0.000408333 | 0.047828399 | **Significant** |
| | | **PEG7.32** | 0.0089 | 7.41667E-05 | 0.008860839 | **Significant** |
| | | **PEG4.27** | 0.0091 | 7.58333E-05 | 0.009059062 | **Significant** |
| **Sorbitol** | | **SorbiWT** | 0.0171 | 0.0001425 | 0.016955823 | **Significant** |
| | | **Sorbi7.32** | 0.0181 | 0.000150833 | 0.01793852 | **Significant** |
| | | **Sorbi4.27** | 0.0202 | 0.000168333 | 0.019999013 | **Significant** |
| **Mannitol** | | **mannWT** | 0.0186 | 0.000155 | 0.018429503 | **Significant** |
| | | **mann7.32** | 0.0189 | 0.0001575 | 0.018723976 | **Significant** |
| | | **mann4.27** | 0.024 | 0.0002 | 0.023716634 | **Significant** |
| **Untreated** | **AtCSD1** | **UntWT** | 0.031 | 0.000258333 | 0.03052831 | **Significant** |
| | | **Unt7.32** | 0.032 | 0.000266667 | 0.031497551 | **Significant** |
| | | **Unt4.27** | 0.0327 | 0.0002725 | 0.032175448 | **Significant** |
| **NaCl** | | **NaCl Wt** | 0.04 | 0.000333333 | 0.039216968 | **Significant** |
| | | **NaCl7.32** | 0.0424 | 0.000353333 | 0.041520872 | **Significant** |
| | | **NaCl4.27** | 0.046 | 0.000383333 | 0.04496646 | **Significant** |
| **PEG** | | **PEGWT** | 0.071 | 0.000591667 | 0.06855768 | **Significant** |
| | | **PEG7.32** | 0.046 | 0.000383333 | 0.04496646 | **Significant** |
| | | **PEG4.27** | 0.045 | 0.000375 | 0.044010586 | **Significant** |
| **Sorbitol** | | **SorbiWT** | 0.09 | 0.00075 | 0.086099675 | **Significant** |
| | | **Sorbi7.32** | 0.018 | 0.00015 | 0.017840294 | **Significant** |
| | | **Sorbi4.27** | 0.022 | 0.000183333 | 0.021761738 | **Significant** |
| **Mannitol** | | **mannWT** | 0.026 | 0.000216667 | 0.025667655 | **Significant** |
| | | **mann7.32** | 0.024 | 0.0002 | 0.023716634 | **Significant** |
| | | **mann4.27** | 0.035 | 0.000291667 | 0.034399513 | **Significant** |
| **Untreated** | **AtCATAL** | **UntWT** | 0.053 | 0.000441667 | 0.051631091 | **Significant** |

| | | | | | | |
|---|---|---|---|---|---|---|
| | **ASE** | **Unt7.32** | 0.065 | 0.000541667 | 0.062949039 | **Non-Significant** |
| | | **Unt4.27** | 0.0074 | 0.000616667 | 0.0071349504 | **Significant** |
| **NaCl** | | **NaCl Wt** | 0.0039 | 0.0000325 | 0.003892468 | **Significant** |
| | | **NaCl7.32** | 0.0059 | 0.000491667 | 0.0057306908 | **Significant** |
| | | **NaCl4.27** | 0.0089 | 7.41667E-05 | 0.008860839 | **Significant** |
| **PEG** | | **PEGWT** | 0.009 | 0.000075 | 0.008959956 | **Significant** |
| | | **PEG7.32** | 0.0107 | 8.91667E-05 | 0.010643431 | **Significant** |
| | | **PEG4.27** | 0.0172 | 0.000143333 | 0.017054136 | **Significant** |
| **Sorbitol** | | **SorbiWT** | 0.0181 | 0.000150833 | 0.01793852 | **Significant** |
| | | **Sorbi7.32** | 0.0027 | 0.0000225 | 0.002696389 | **Significant** |
| | | **Sorbi4.27** | 0.0646 | 0.000538333 | 0.062573941 | **Non-Significant** |
| **Mannitol** | | **mannWT** | 0.015 | 0.000125 | 0.014888984 | **Significant** |
| | | **mann7.32** | 0.023 | 0.000191667 | 0.022739671 | **Significant** |
| | | **mann4.27** | 0.043 | 0.000358333 | 0.042095992 | **Significant** |
| **Untreated** | **AtERF5** | **UntWT** | 0.009 | 0.000075 | 0.008959956 | **Significant** |
| | | **Unt7.32** | 0.0016 | 1.33333E-05 | 0.001598731 | **Significant** |
| | | **Unt4.27** | 0.0019 | 1.58333E-05 | 0.001898211 | **Significant** |
| **NaCl** | | **NaCl Wt** | 0.0034 | 2.83333E-05 | 0.003394275 | **Significant** |
| | | **NaCl7.32** | 0.0081 | 0.000675 | 0.007783153 | **Significant** |
| | | **NaCl4.27** | 0.055 | 0.000458333 | 0.053526785 | **Significant** |
| **PEG** | | **PEGWT** | 0.056 | 0.000466667 | 0.054473223 | **Significant** |
| | | **PEG7.32** | 0.0038 | 3.16667E-05 | 0.003792849 | **Significant** |
| | | **PEG4.27** | 0.0044 | 3.66667E-05 | 0.004390414 | **Significant** |
| **Sorbitol** | | **SorbiWT** | 0.085 | 0.000708333 | 0.081515379 | **Non-Significant** |
| | | **Sorbi7.32** | 0.005 | 4.16667E-05 | 0.004987624 | **Significant** |
| | | **Sorbi4.27** | 0.0055 | 0.000458333 | 0.0053526785 | **Significant** |
| **Mannitol** | | **mannWT** | 0.016 | 0.0005 | 0.0158249597 | **Significant** |
| | | **mann7.32** | 0.0045 | 0.0000375 | 0.004489974 | **Significant** |
| | | **mann4.27** | 0.055 | 0.000458333 | 0.053526785 | **Significant** |
